# Insights into non-crossover recombination from long-read sperm sequencing

**DOI:** 10.1101/2024.07.05.602249

**Authors:** Regev Schweiger, Sangjin Lee, Chenxi Zhou, Tsun-Po Yang, Katie Smith, Stacy Li, Rashesh Sanghvi, Matthew Neville, Emily Mitchell, Ayrun Nessa, Sam Wadge, Kerrin S Small, Peter J Campbell, Peter H Sudmant, Raheleh Rahbari, Richard Durbin

## Abstract

Meiotic recombination is a fundamental process that generates genetic diversity by creating new combinations of existing alleles. Although human crossovers have been studied at the pedigree, population and single-cell level, the more frequent non-crossover events that lead to gene conversion are harder to study, particularly at the individual level. Here we show that single high-fidelity long sequencing reads from sperm can capture both crossovers and non-crossovers, allowing effectively arbitrary sample sizes for analysis from one male. Using fifteen sperm samples from thirteen donors we demonstrate variation between and within donors for the rates of different types of recombination. Intriguingly, we observe a tendency for non-crossover gene conversions to occur upstream of nearby PRDM9 binding sites, whereas crossover locations have a slight downstream bias. We further provide evidence for two distinct non-crossover processes. One gives rise to the vast majority of non-crossovers with mean conversion tract length under 50bp, which we suggest is an outcome of standard PRDM9-induced meiotic recombination. In contrast ∼2% of non-crossovers have much longer mean tract length, and potentially originate from the same process as complex events with more than two haplotype switches, which is not associated with PRDM9 binding sites and is also seen in somatic cells.

## Main text

Meiotic recombination shapes evolution by generating new combinations of alleles, which serve as a foundation for adaptation and speciation through natural selection. The accurate execution of recombination is essential for maintaining genome integrity through meiosis and for fertility in many organisms^1^. In meiotic homologous recombination, homologue-templated repair of programmed DNA double-strand breaks (DSBs) causes the exchange of genetic material between homologous chromosomes. In humans and many other vertebrates meiotic DSB generation is localised by sequence-specific binding of the zinc-finger PRDM9 protein^2^. A minority of DSBs resolve into crossovers (COs), which result in reciprocal exchange of alleles between homologues over a large chromosomal distance. Conversely, many more DSBs resolve into non-crossovers (NCOs), characterised by a unidirectional transfer of genetic information over short intervals (gene conversion). Additionally, complex recombination events, subclassified into complex COs and complex NCOs, have been observed, albeit in small numbers^3–5^. In complex COs, one or more NCO tracts are present near the CO breakpoint. In complex NCOs, multiple NCO tracts cluster in close proximity. Despite the key role of recombination in genetic inheritance, our knowledge and understanding of these processes remains incomplete, particularly in humans.

First, significant gaps exist in our knowledge about NCO events. While the relative frequency of NCOs has been estimated to be an order of magnitude higher than that of COs^4, 6, 7^, we lack a clear understanding of the variation in NCO rates across different genetic hotspots and across individuals. Additionally, there is inconsistency in the literature about the length distribution of NCO tracts, with estimated mean values varying from low tens^8, 9^ to hundreds of bp^4, 6, 10^. This stems from the fact that NCO events are difficult to detect: their detection relies on the conversion of pre-existing heterozygous markers, so the resolution of direct measurement of NCO events depends on the local density of heterozygous variants.

Our understanding of complex events is even more limited. It is thought that complex events are the result of patchy mismatch repair of the heteroduplex DNA in recombination^3^. Complex COs have been observed at rates of ∼0.3% in males and more frequent in females (∼1.3%), increasing with maternal age^3, 5^. NCO tracts involved in complex COs were generally longer than tracts in simple NCOs, suggesting a separate pathway. Complex NCOs have also been observed^4, 5^, thought to be a product of alternating mismatch strand preference. This is supported by evidence of GC-bias in both converted and unconverted markers, which is a signature of mismatch repair^11^.

Maps of population-averaged variation in crossover rate across the genome and many other properties of recombination have been obtained using pedigrees and statistical analysis of population sequencing data. While pedigree sequence data^4, 5, 8, 12, 13^ allows high quality variant calling and phasing, and hence reliable calling of CO and NCO events, these studies generate relatively few recombination events from large numbers of individual genome sequences, with each person contributing only a limited number of recombinations. Statistical approaches exploit the accumulation of recombination events across generations and can provide fine-scale population-wide averages of crossover rates^14–16^, but they rely on assumptions such as simplified models of population history, a constant mutation rate and selective neutrality. Furthermore, statistical analysis of NCOs based on comparison of long identity by descent segments^17, 18^ is also vulnerable to sequencing errors.

An alternative approach involves direct sequencing of many sperm cells from a single individual, each almost certainly representing an independent meiosis. Early studies used long-range PCR amplification of selected hotspots but were limited to a few loci. Single-cell methods allow insights into COs genome-wide but lack resolution for NCOs^19, 20^. Recently, advances in long-read high-throughput sequencing have allowed sperm typing using a shotgun approach^21^, giving information about which alleles co-occur on the same parental chromosome since each read typically spans multiple markers^22–24^. Despite this advance, long-read sequencing has so far not been used to study NCOs, likely due to high error levels in earlier long-read technology confounding converted markers with errors.

Here we show that long-read sperm sequencing with Pacific Biosciences circular consensus sequencing (CCS) technology overcomes previous limitations, due to its accuracy removing the confounding of gene conversion with error. In particular, it allows the detection of both CO and NCO recombination events from a single sperm sample, without requiring additional data such as pedigree or trio sequencing. We show previously unreported variation between individuals in patterns of recombination and their molecular signatures. Using likelihood-based inference, we estimate the tract length distributions for a mixture of two distinct NCO recombination processes. Intriguingly, we find that COs tend to be located on one side of nearby PRDM9 motifs, while NCOs are on the opposite side.

## Results

We obtained 15 sperm samples from 13 donors, with ages at donation varying between 24 and 74, together with two blood samples as controls, one from neonatal cord blood and one from an 82-year old female (Tables 1, S1, see Methods for details). Eleven individuals donated one sperm sample, two of whom are monozygotic twins, and two individuals donated two sperm samples each, at different ages. We assigned identifiers to the sperm donors based on their *PRDM9* genotype. Ten samples from nine donors, including the two twins, were homozygous for the PRDM9_A_ allele (samples AA1-s1, AA-s2, AA2-t1, AA2-t2, and AA3 to AA9), which is the most prevalent in European ancestry^25^. One donor (AB) was heterozygous for the PRDM9_A_ allele and the PRDM9_B_ allele, and one donor (AD) for PRDM9_A_ and the PRDM9_D_ allele. Interestingly, PRDM9_D_ is reported to be a de-stabilizing allele that appears to cause elevated variation of the zinc-finger array in sperm^26^, and was previously exclusively found in the Finnish population^25^. One donor (with two samples AN-s1 and AN-s2) was heterozygous to PRDM9_A_ and to a novel PRDM9 allele not reported previously. The zinc-finger array of this new allele is derived from PRDM9_A_, with an additional repeat of the second complete zinc finger (we supply the new sequence in the Supplementary Text). The two control samples are labelled C1 and C2.

Nine sperm samples from the TwinsUK^27^ project (hereafter “TwinsUK samples”) and the two control samples were sequenced on a Pacific Biosciences Sequel II instrument, and six sperm samples from the Sudmant lab (hereafter “SL samples”) on a Pacific Biosciences Revio instrument. The Revio has lower overall error rates, but also lower precision of its most confident base calls. We obtained nearly 35x average CCS sequence coverage per sperm sample (range 16x-57x), with average read length 21,778 bp and a total of 101,486,297 reads.

### Identification of candidate recombinant reads

To ensure accurate read mapping and avoid reference bias we assembled a haplotype-resolved de-novo assembly for each individual using `hifiasm`^28^, resulting in two sets of chromosome sequences, which we refer to below as “haplotypes” (Fig. 1A). We then separately aligned each read to each of its donor’s two haplotypes, retaining for further analysis only 74,380,961 reads that map unambiguously to both haplotypes and satisfy other quality control filters (see Methods).

**Fig. 1.**
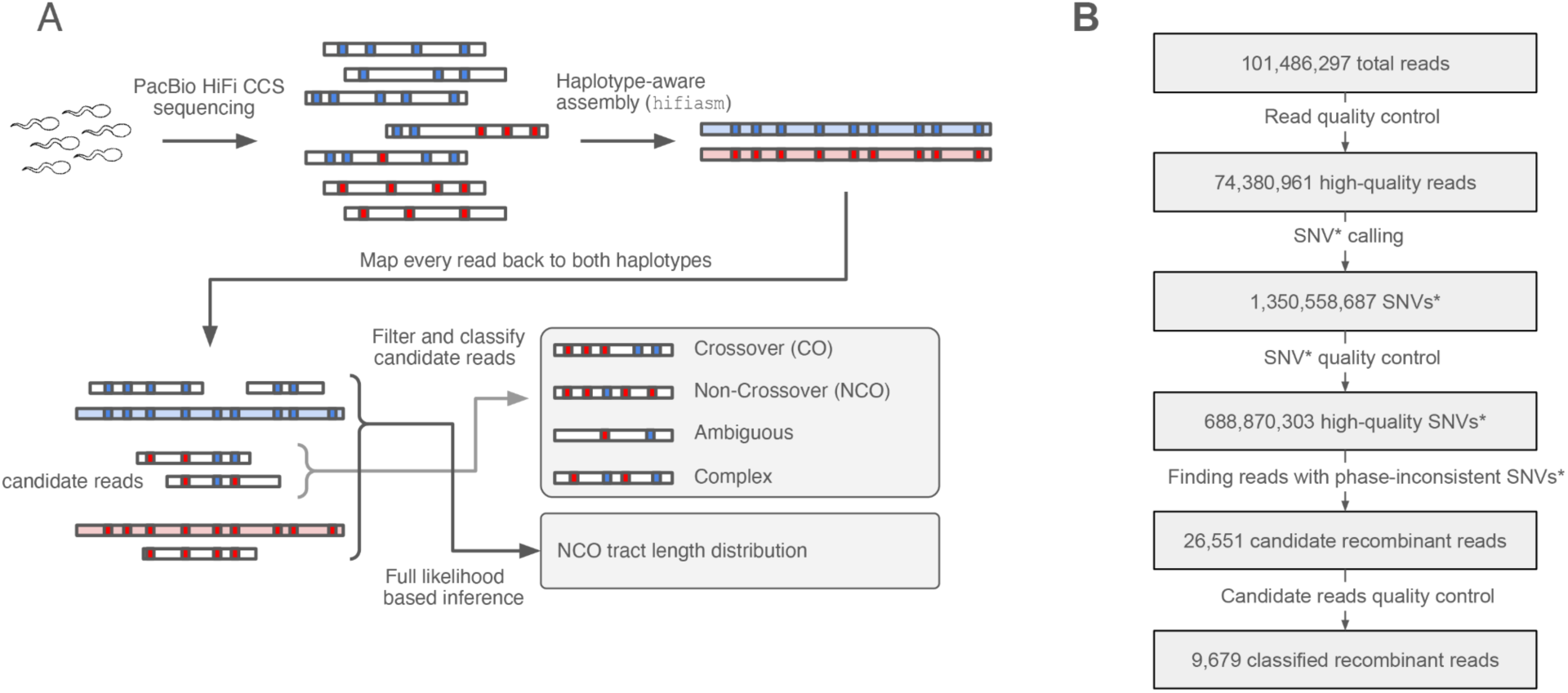
(A) workflow for data processing. (B) numbers as the entire data set passes through the workflow. SNVs* in boxes 3 and 4 indicates candidate SNVs assigned at read level not at sample level.

We next generated candidate single nucleotide variant (SNV) calls on each read separately, by retaining only sites that matched to one haplotype but not the other. To remove false positives, we filtered out candidate SNVs that occur in regions of low complexity, tandem repeats or that mapped to haplotype regions where few other reads aligned (see Methods). Additionally, a key challenge in calling recombination events is distinguishing between sequencing errors and a true NCO converting a single marker. We controlled for this by removing candidate SNVs with base quality (BQ) on their parent read below a threshold, and further quality controls detailed in Methods. This resulted in 688,870,303 SNVs across all reads, with a mean 9.85 SNVs per read (Extended Data Fig. 1).

By construction each allele matches one of the two haplotypes. A candidate recombinant read will have SNVs with alleles from both haplotypes. In order to confirm a haplotype switch between adjacent SNVs we required that the region between them is continuously spanned by at least 3-fold coverage on each haplotype, to be confident that the switch is not an artefact of assembly. Finally, we removed candidates which share a switch between the same two SNVs with another read. Such repeated switches likely indicate an assembly or phasing error, since at our read coverage the chance of two independent recombination events between any pair of SNVs is extremely low, even in hotspots.

Because we only observe a few markers per read, we may not be able to confidently classify an observed pattern of haplotype switches in a read to a CO or NCO. For quantitative inference of NCO properties, we use a statistical likelihood-based method as described below, but for qualitative analysis we carried out an initial classification of candidate reads. We label a candidate read as *CO* if it contains a single switch with at least two SNPs supporting it on each side (e.g. two SNPs from haplotype 1 to the left and two SNPs from haplotype 2 to the right); *NCO* if there are two switches; *complex* if it has three switches or more; or *ambiguous* if there is only one switch but not enough SNVs supporting it (e.g. only SNP on one side or the other). For this analysis we added back some SNVs not used during candidate read generation to improve resolution (see Methods for details). A substantial fraction of the candidate recombinant reads were checked manually in a read alignment viewer presenting the aligned reads to both haplotypes, resulting in some tuning of parameters and the removal of 13 NCO events (see Methods for details). Ultimately we detected 4,460 COs, 1,759 NCOs, 351 complex and 3,109 ambiguous events across all sperm samples (Table 1, Fig. 2A, Table S1). The number of switches in complex events varies between 3 and 15, with 3 and 4 being most frequent (Extended Data Fig. 2).

**Fig. 2.**
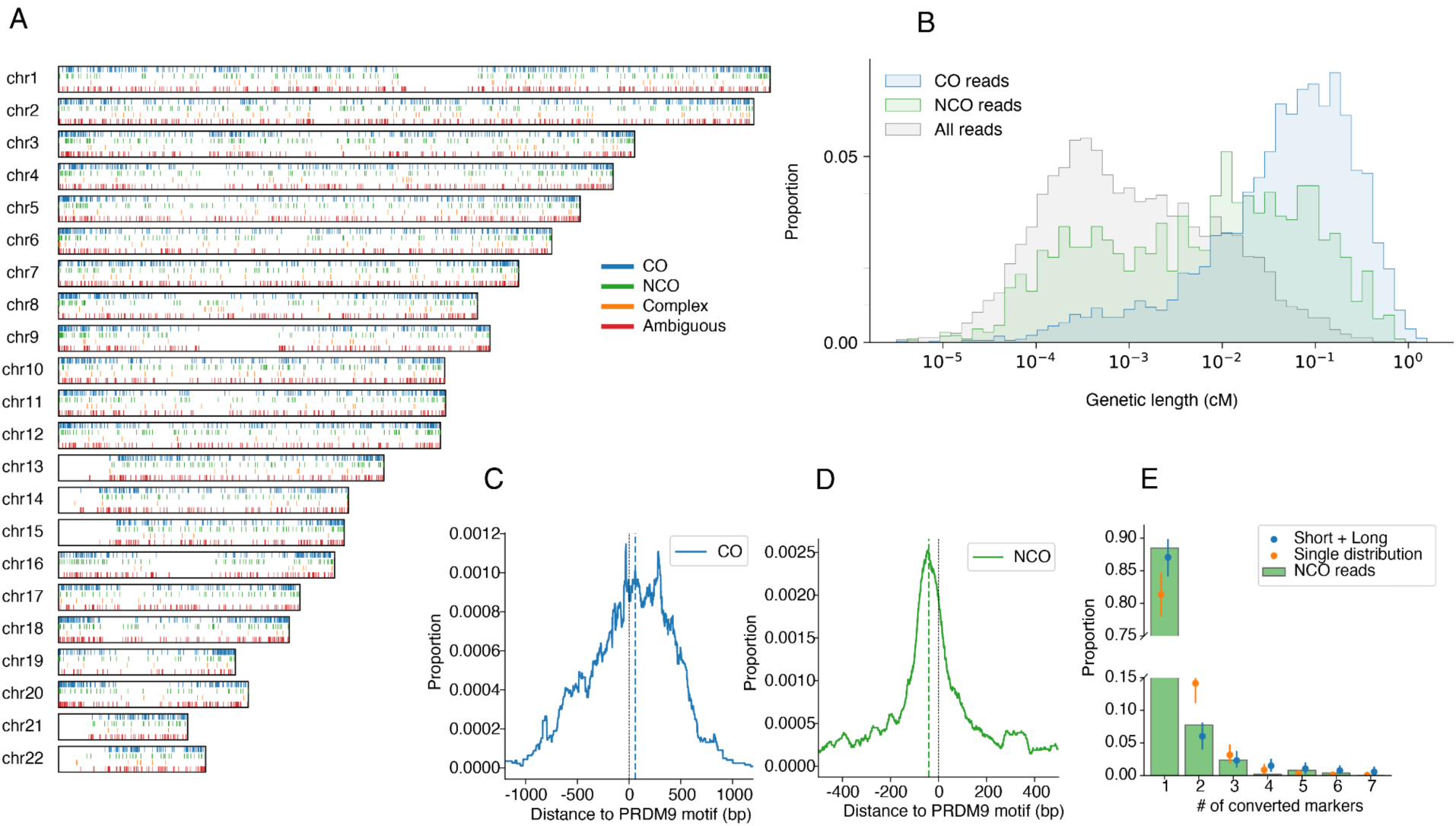
(A) genomic distribution of called events. (B) enrichment of CO and NCO reads in regions of high recombination rate according to the male genetic map. (C) accumulated weight of crossover intervals for called COs relative to the closest oriented DSB-PRDM9 motif; the dashed line indicates the median (D) relative position of converted SNV for called NCOs relative to the closest oriented DSB-PRDM9 motif. (E) fit of observed number of consecutive converted SNVs in NCO reads to single geometric and mixture of short and long geometric tract length distributions.

**Table 1.** Details of samples. The number of classified recombinant reads with age, sequencing coverage, CO, NCO, ambiguous or complex recombination events, per sample. Sample pairs AA1-s1/AA1-s2 and AN-s1/AN-s2 are from the same donor at different ages. Samples AA2-t1 and AA2-t2 are monozygotic twins. C1 and C2 are blood samples. Rows shaded in grey indicate samples sequenced on Revio machines whereas unshaded rows contain samples sequenced on Sequel II machines.

| Sample | Age | Coverage | CO | NCO | Ambiguous | Complex |
| --- | --- | --- | --- | --- | --- | --- |
| AA1-s1 | 62 | 20.0x | 161 | 45 | 95 | 3 |
| AA1-s2 | 74 | 16.7x | 130 | 31 | 78 | 8 |
| AA2-t1 | 24 | 31.3x | 262 | 72 | 135 | 45 |
| AA2-t2 | 36 | 24.2x | 179 | 82 | 110 | 54 |
| AA3 | 74 | 31.2x | 205 | 73 | 126 | 31 |
| AA4 | 47 | 21.5x | 180 | 51 | 80 | 24 |
| AA5 | 62 | 53.6x | 520 | 233 | 419 | 13 |
| AA6 | 31 | 51.8x | 447 | 212 | 326 | 11 |
| AA7 | 32 | 57.0x | 501 | 212 | 346 | 18 |
| AA8 | 31 | 49.8x | 442 | 171 | 346 | 12 |
| AA9 | 28 | 48.8x | 425 | 181 | 346 | 9 |
| AN-s1 | 27 | 31.2x | 233 | 91 | 161 | 62 |
| AN-s2 | 40 | 24.3x | 202 | 71 | 118 | 33 |
| AB | 33 | 40.3x | 363 | 149 | 304 | 9 |
| AD | 26 | 23.7x | 210 | 85 | 119 | 19 |
| **Totals** |  | **525.5x** | **4,460** | **1,759** | **3,109** | **351** |
| C1 | 0 | 70.2x | 2 | 20 | 7 | 22 |
| C2 | 82 | 30.1x | 2 | 19 | 3 | 2 |

It is apparent from Table 1 that there are systematic differences between callsets from Revio machines and Sequel II machines, with a higher rate of NCO and ambiguous candidate reads in the Revio data (CO:NCO Chi-squared P=9e-5). We attribute this to differences in the base quality binning between platforms, which affects our ability to distinguish NCOs from sequencing errors. Therefore below when we compare individuals we stratify by callset.

Processing the control samples (both sequenced on Sequel II) in identical fashion, we detected 4 COs, 39 NCOs, 24 complex and 10 ambiguous events. After correcting for coverage these represent 0.5%, 11.6%, 35.8% and 1.7% respectively of the rates seen in sperm samples. At worst these bound the error rate for each category in the TwinsUK sperm callset, which was also called from Sequel II data. However, almost all the NCO and complex control events looked real in manual checks, and the other potential explanation for them is that they represent the products of somatic rather than meiotic recombination. Under that interpretation, while the somatic crossover rate is very low compared to the meiotic rate, somatic gene conversion, assigned to NCOs and in particular complex events, is relatively higher, potentially reflecting the consequences of mismatch repair associated with the fixing of somatic double stranded breaks^29^.

### Relationship of recombination events to the genetic map

When combined across all samples, the distribution of detected CO events along each chromosome largely follows the Behrer et al. male-specific genetic map^30^ as expected (Extended Data Fig. 3). Using the map to calculate the genetic length of the CO intervals (the regions in candidate CO reads between the pair of SNVs flanking the crossover), they have an average map recombination rate of 32.3 cM/Mb, compared to a background rate of 1.1 cM/Mb in all reads processed through the same pipeline (Kolmogorov-Smirnoff two-sample test (KS), P<1e-100). In contrast the average map recombination rate at NCO converted markers is 15.1 cM/Mb, again significantly enriched compared to the background rate (KS P<1e-100), but less so than the rate calculated above for COs. This reflects the fact that the Behrer et al. map is calculated from crossovers, and while both COs and NCOs are initiated by DSBs and so reflect the genomic distribution of meiotic DSBs, the relative rates can vary along the genome and perhaps between samples^19^, so the NCO map only approximately follows the CO map.

To confirm differences in the distribution of recombination events controlling for possible ascertainment effects induced by variation in SNV density we compared the distribution of genetic lengths between the second SNV and second-to-last SNV of CO, NCO and all reads (Fig. 2B; see Methods). We find that the distribution of NCO events is significantly different from that of CO events, following the CO map less well (even when considering just the TwinsUK callset), while both are significantly different from the background distribution of all other reads (Fig. 2B; KS, all P<1e-30).

Finally, we considered complex events. The average crossover rate according to the Behrer et al. map for these reads is only 1.3cM/Mb, substantially lower than the values for either COs (9.0cM/Mb; lower than given above because it is across the whole read) or NCOs (4.0cM/Mb) and close to the background rate of 1.1cM/Mb. This suggests that complex recombination events may arise largely through a different process. We note that the complex event reads are not clustered and wherever they are found there are also reads consistent with both haplotypes, supporting that they are not artefacts of local assembly quality.

Mapping of long reads to an individualised assembly allowed us to call CO and NCO events in repeat-rich regions such as the subtelomeric regions, previously inaccessible to short-read based pedigree studies or earlier sperm sequencing studies. An assessment of the genomic distribution of CO and NCO events showed that both are elevated near telomeres, consistent with previous results^5^ (Extended Data Fig. 4). We observed gradual elevation in the number of events at low resolution (10Mbp resolution in 100Mbp nearest to telomeres) and also at higher resolution (1Mb resolution in 20Mb nearest to telomeres), previously seen also in mice^8^. In particular, we observed more NCOs than COs at the 1Mbp region near the telomere, and confirmed that this is not due to an excess of marker density leading to higher NCO detectability. We note an apparent difference in the distribution of NCOs between the TwinsUK and the SL samples, possibly due to the batch effects discussed above.

### Relationship of recombination events to the double strand break map

To better understand the relationship between recombination events and the DSBs that initiate them, we compared CO and NCO regions to a previously published DSB map of 28,286 human recombination hotspots, based on the measurement of the occupancy of a meiotic DSB repair protein (DMC1) in chromatin from testes of a PRDM9 A/A homozygous male^31, 32^. Each called hotspot is centred around a 31bp PRDM9 motif^32^, which we will refer to as DSB-PRDM9 motifs. We found that 70% of CO reads and 47% of NCO reads overlapped a DSB-PRDM9 motif (Fisher’s exact test for CO vs. NCO, P=1.2e-8), compared with 15% of all reads (Fisher’s exact test, P<1e-100); we note that reads may contain PRDM9 motifs not in our DSB hotspot list. In 57% of the CO reads overlapping a DSB-PRDM9 motif the motif was contained within the CO interval, while in 59% of NCO reads overlapping a DSB-PRDM9 motif the motif was contained within the interval spanned by the two markers surrounding the NCO tract (not significantly different). In contrast to both COs and NCOs, only 15% of complex event reads overlap a DSB-PRDM9 motif, the same fraction as for all reads.

The localization of PRDM9 motifs within DSB hotspots allows us to study the relative positions of recombination events and the PRDM9 binding site. Focusing on CO reads containing a DSB-PRDM9 motif, and with CO intervals shorter than 500bp to increase the resolution of our analysis, we calculated the distribution of the offset from the motif centre to each base in the CO interval, weighting each observed distance inversely to interval length. The resulting distribution has an average distance of ∼380bp to the motif centre (Fig. 2C); we note that this may be inflated by using even weights to handle the uncertainty in position of the crossover within the CO interval. Strikingly, we observed an asymmetry in distances, with 58% of the interval weight downstream of the DSB-PRDM9 motif, and with a median around ∼59bp (Fig. 2C, permutation test P=0.04).

We similarly estimated the distribution of offsets between converted markers in NCO reads and the DSB-PRDM9 motif. Because NCO tracts are typically short (<100bp, see below), focusing on converted markers allowed us to localise the recombination breakpoints with higher resolution than for COs. The resulting distribution showed an average distance of ∼280bp to the motif centre (Fig. 2D). We also observed an asymmetry here, but in the other direction, with 60% of the converted bases occurring upstream of the DSB-PRDM9 motif, and with a median around ∼-38bp (Fig. 2D, permutation test P=1e-4). Together, these results suggest a directional influence of the PRDM9 binding site on the location of CO and NCO events.

### Inference of NCO tract lengths

NCO tracts cannot be observed directly from the sequencing data: they only leave evidence at heterozygous markers that are converted. Most tracts convert only a single SNP (1,524 of 1,642 candidate reads; 92.8%), although there is one that converts 7 SNPs (Extended Data Fig. 2). We therefore need a statistical inference approach to infer the distribution of tract lengths.

As in previous work^8, 33–35^ we model the tract length as following a geometric distribution, which is natural for a repair process that proceeds along the chromosome with a constant probability of ending at each base. However, instead of fitting just to the candidate NCO reads we fit a full likelihood model across a much larger set of reads that were not ascertained on whether or not they show signs of recombination, allowing for unobserved events. In practice we use all TwinsUK reads with at least three SNVs that satisfy quality control criteria; we restrict to TwinsUK samples to avoid batch effects and the higher NCO false positive rate in SL data. When we fit a model with a single geometric distribution to this data (allowing a nuisance parameter *q* to fit the fraction of recombination events resulting in a CO), we inferred a mean tract length of 217.2bp. This is long compared to recent estimates, and furthermore, the expected distribution of the number of converted markers under this inferred distribution is inconsistent with the observed data (Fig. 2E).

Previous work has also observed a small fraction of NCO tract to be especially long^4, 5^, thought to arise from a separate pathway or to be a part of a complex recombination event. Indeed, in a recent pedigree study of baboons^12^ a non-negligible fraction of long (>1kbp) NCO tracts was also observed, and a mixture of two geometric distributions (one short with mean ∼24bp and one long with mean >4kbp) was fit to the data. Consequently, we extended our model to fit the tract length distribution as a mixture of two geometric distributions, now estimating the mixture proportion *m* of the long component, and the mean tract lengths of the two geometric distributions *L_1_* and *L_2_*. With this model we inferred a mixture proportion of 1.8% (95% nonparametric bootstrap confidence interval CI: 1.1%-2.9%), with the major component capturing short tracts with inferred mean L_1_=34.6bp (CI: 23.5-57.54 bp) and a minor component modelling longer tracts with mean L_2_=7,214.1bp (CI: 1,022.9-11,306.2 bp). We note in particular the high uncertainty of the long tract distribution. The log-likelihood of the mixture distribution is -26726.4, a greater than 10^59^-fold improvement over the value of -26863.9 for the single component model, resulting in a very highly significant Akaike Information Criterion (AIC) value (1.3e-59 relative likelihood). Moreover, we observe a substantially better fit for the expected distribution of converted markers (Fig. 2E). With these parameters, we estimate that only ∼1.4% of NCO tracts will be observed, and that 14.2% (CI: 10.5%-20.2%) of recombinations result in crossovers, although we note that this fraction can be used by the model to accommodate systematic effects, so is potentially subject to bias. However, while only approximately 2% of NCO events come from the longer component, we estimate that 18% of detected NCOs are caused by long events.

### GC-bias in NCO tracts

GC-biased gene conversion (gBGC) in NCO tracts is the preferential incorporation of G or C nucleotides during mismatch repair in non-crossover events. We observed a GC-bias of 58.3% (CI: 55.8%-60.7%) in 1,516 converted SNPs in 1,385 NCO events. We see no significant difference in multi-SNP tracts (58.4%, CI: 51.7%-64.8%). This is towards the lower end of the Haldorsson et al.^5^ estimate for NCOs in males (57.1%–69.1%). When we limit our analysis to NCO reads overlapping a PRDM9 motif within the DSB map, we do observe a significantly higher value of 63.6% (CI: 59.8%-67%, Fisher’s exact P=0.00013).

### Individual variation in CO and NCO events

A key advantage of our study design is that we can observe large numbers of recombination events per sample, proportional to read coverage under experimental control rather than the number of offspring of an individual. Increased coverage gives greater power to investigate variability between individuals or even between samples from the same individual.

To this end, we first examined the variability in genetic lengths in centimorgans (cM) of CO reads, which reflect the degree COs follow the CO recombination map (Fig. 3A). The AD individual had significantly smaller cM than other samples (see Methods; Anderson-Darling statistic with permutation testing, P<1e-4), consistent with it departing from the CO map determined largely by the European-dominant PRDM9A allele (Fig. 3A and Extended Data Fig. 5A). The two samples from AN (AN-s1/s2) and the sample from AB were significantly differently distributed than the rest of the samples, even when excluding the AD individual from the comparison (permutation tests P<1e-4 and P=0.004 respectively, Fig. 3A and Extended Data Fig. 5B and 5C), again consistent with these individuals carrying a different PRDM9 allele. Finally, within AA samples, the two samples from AA1 were statistically differently distributed to all other AA samples, following the CO map more closely on average than others (permutation tests P=0.0003 and P=0.0013, Fig. 3A and Extended Data Fig. 5D).

**Fig. 3.**
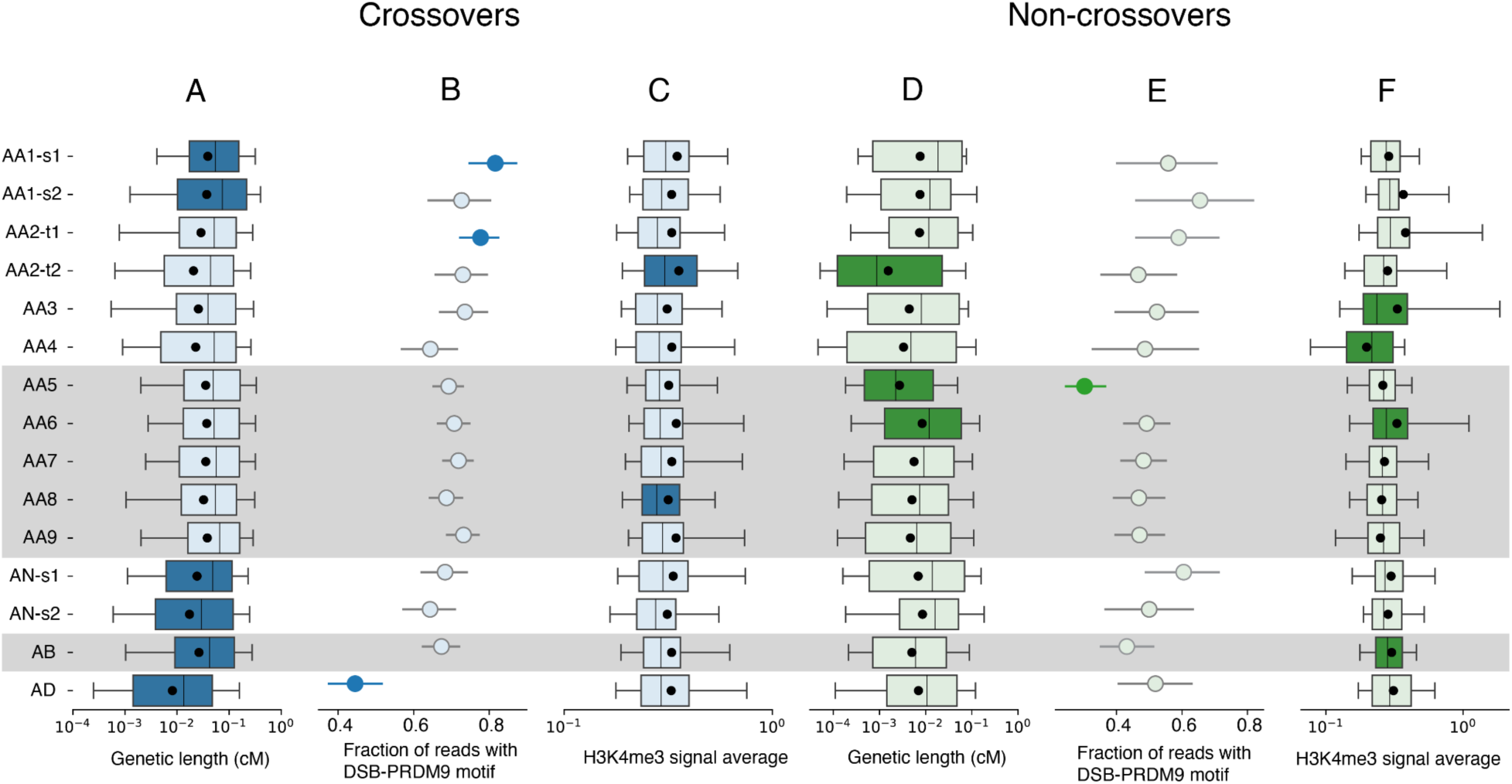
Individual variation in properties of meiotic recombination, per-sample: (A) the distribution of genetic length in cM of CO reads. (B) the fraction of CO reads that overlap a DSB-PRDM9 motif. (C) the distribution of the average H3K4me3 signal strength across CO reads. (D) the distribution of genetic length in cM of NCO reads. (E) the fraction of NCO reads that overlap a DSB-PRDM9 motif. (F) the distribution of the average H3K4me3 signal strength across NCO reads. In (A), (C), (D) and (F) boxes show the 25%, 50% and 75% quantiles, whiskers indicate 10% and 90% quantiles, and the black circle indicates the mean. In (B) and (E) the lines indicate the 95% binomial confidence interval. Coloured boxes and circles indicate samples that are statistically significantly different from other samples. Shading indicates SL samples sequenced with the Revio machine.

A comparison with the DSB hotspot map allows for an additional perspective on individual variation. We first compared the fraction of reads that overlap a PRDM9 motif associated with a DSB hotspot in the Hinch et al. map^32^ (Fig. 3B). In accordance with the disparity in genetic length distribution, only 44.5% of reads from the AD sample overlap a motif, significantly lower than the other samples (Fisher’s exact test, P=1.8e-13). Excluding AD, sample AA1-s1 (but not AA1-s2, an older sample from the same individual) has a significantly larger proportion of reads overlapping a motif (81.5%, Fisher’s exact P=0.002), in line with it following the CO map more closely than other AA samples. Sample AA2-t1 (the twin sample obtained at a younger age; but not AA2-t2) also has a significantly larger fraction (77.6%, P=0.009 excluding AD and AA1-s1). Except for these three samples, we detected no other significant pairwise differences in CO reads.

PRDM9 facilitates recombination by acting as a histone H3K4 trimethyltransferase. We compared CO reads with a dataset of H3K4me3 ChIP-Seq peak calls derived from testis tissue by the ENCODE consortium^36^ (see Methods). DSB hotspots have been shown to overlap with H3K4me3 marks in testis^31^. We calculated the average signal across the detectable interval of each read, and compared the distribution of this between reads from different samples (Fig. 3C). Sample AA2-t2 (the twin sample obtained at an older age), but not AA2-t1, has statistically significantly higher signal on average to the others (permutation test P<1e-3, Extended Data Fig. 6A). Excluding AA2-t2, sample AA8 has slightly, but significantly, lower H3K4me3 signal than all other remaining samples (permutation test P=0.017, Extended Data Fig. 6B).

We next turned to analyse differences between samples in the genomic distribution of NCO events, by comparing the distribution of the genetic lengths of NCO reads between samples as we did for CO reads (Fig. 3D). For these comparisons we stratified the samples by sequencing platform, for reasons described above. For the SL samples sequenced on Revio, AA5 differs substantially, with significantly lower average cM read length (permutation test P<1e-4, Extended Data Fig. 7A). Removing AA5 from the comparison, sample AA6 has marginally higher cM read length (permutation test P=0.004, Extended Data Fig. 7B). Amongst the TwinsUK samples sequenced on Sequel sample AA2-t2 (but not AA2-t1) has significantly lower cM read length (permutation test P<1e-4, Fig. 3B and Extended Data Fig. 7C). The proportion of NCO reads overlapping DSB-associated PRMD9 motifs is also significantly lower for AA5, both when stratifying by callset (Fig. 3E) and compared to all other samples. (P<1e-20 using Fisher’s method to combine pairwise Fisher’s exact tests). After removing AA5 no other samples are significant.

We also compared the H3K4me3 signal across NCO reads (Fig. 3F). Amongst the TwinsUK samples, AA4 was substantially lower than other samples (permutation test P<1e-4, Extended Data Fig. 8A). Removing AA4, AA3 was significantly higher than remaining TwinsUK samples (permutation test P=0.002, Extended Data Fig. 8B). For the SL samples AA6 was significantly higher than other samples (permutation test P=0.001, Extended Data Fig. 8C). Removing AA6, AB is significantly higher than the rest of other samples (P=0.007, Extended Data Fig. 8D).

Finally, there is clearly substantial individual variation in the rate of complex events compared to the sum of COs and NCOs (Table 1; P=2e-74 chi-squared test). There is no significant correlation of this ratio to age (R^2^=0.02).

In summary, we see many differences between samples in the properties of both COs and NCOs. Some of these are consistent with the *PRDM9* genotype as expected, but others involve differences between individuals with the same genotype, and in some cases between repeat samples from the same individual or between monozygotic brothers. Further, differences in CO properties are not consistent with differences in NCO properties; it appears that the relative patterns of NCOs to COs are also variable between samples.

## Discussion

We have shown here a new approach to using long high accuracy shotgun sequencing reads from sperm samples to study recombination processes. In particular, the very high accuracy of Pacific Biosciences CCS base calls allows us to accurately identify non-crossover events, extending beyond previous genome-wide sperm sequencing studies that were limited to crossover events^19–21^. Using this approach we have been able to characterise individual variation in NCO properties for the first time, showing that this does not mirror individual variation in CO properties that had been studied before. In several cases we see differences between samples taken from the same individual at different times (approximately 12 years apart) or monozygotic twins, suggesting that stochastic or environmental factors play a significant role in meiosis outcomes.

We provide clear evidence that NCOs are not well fit by a single geometric distribution, with a much better fit by a mixture of two distributions composed of a short component of mean length under 50bp (we estimated 34bp) accounting for approximately 98% of events and a longer component allowing for tracts of length multiple kilobases accounting for the remaining events. We do not have enough data to distinguish whether the larger component is itself a mixture. However we note that, as others have observed, we see complex events extending over many SNPs, and suggest that the longer NCO component we detect may come from the two-switch fraction of these complex events, or indeed from longer complex events that extend beyond the end of a read leaving only two switches within the read. Moreover, our observations that the complex events are not associated with PRDM9 sites, and that they are also found relatively frequently in control blood DNA suggests that they, and the longer NCO component, may arise from a recombination process not associated with PRDM9-mediated DSBs, which is also active in somatic cells. Even in sperm we expect to see the results of somatic/mitotic as well as meiotic processes. Although they are only about 2% of events, the long category of NCOs typically span heterozygous sites and account for a substantial fraction of the converted SNVs. The potential lack of association between these long-component converted SNVs and the PRDM9-mediated process would in part explain the reduction in association of NCOs compared to COs with DSB-PRDM9 sites and the male genetic map.

The observation of opposite tendencies for NCOs and COs to be found upstream and downstream of PRDM9 motifs respectively is surprising. Previous studies have established that PRDM9 acts as a chromatin remodeler as well as trimethylating lysines on the histone H3 tail, thus influencing where the DSB required for DNA exchange between chromosomes is formed, although this process was believed to be symmetric^37^. Further, PRDM9 remains bound to DNA through subsequent steps in meiosis, interacting with other recombination proteins through its KRAB domain^38^. Although these steps provide the potential for an influence of PRDM9 orientation on recombination outcome, how that might be achieved is an open question.

The experimental simplicity of using a single sperm sample allows this approach to be used on other species for which sperm, or in some cases very large quantities of eggs, can be obtained. It could also be applied to study the effects on recombination outcomes of mutations in genes known to be involved in recombination, by sequencing sperm from males carrying the mutations. In humans and most other species a limitation is that the approach can only be used to study male meiosis. Other limitations are that long-range interactions such as crossover interference cannot be studied, and the full extent of complex events, including whether they resolve to COs or NCOs, cannot be reliably determined.

Finally, we can consider the possibility of much higher sequencing depths of perhaps thousands of fold coverage on just one or a few samples. In principle this would allow direct study of repeated recombination events in the same sample at individual hotspots and the study of rare events such as non-allelic homologous recombination (NAHR) and recombination-associated mutagenesis^32^. This advance would offer deeper insights into the mechanisms driving meiotic recombination and their implications for genetic diversity and evolution.

## Methods

### Sperm sample extraction and sequencing

We obtained 15 sperm samples from 13 donors of European ancestry. Eleven individuals donated one sample, and two individuals donated two samples at different ages (AA1-s1 and AA1-s2; AN-s1 and AN-s2), with ages at donation varying between 24 and 74 (see Supplementary Table 1). Samples AA2-t1 and AA2-t2 are monozygotic twins.

The TwinsUK cohort consists of more than 14,000 volunteers, mostly middle-aged females, who have participated over the last 30 years in a longitudinal cohort study. This has included lifestyle and health questionnaires, biomedical measurements, biological sample collection, and the generation of multi-omics profiles (such as genetics, metagenomics, and metabolomics), over multiple visits.

For the Twins UK samples, we obtained high molecular weight (HMW) DNA from bulk sperm samples using the Circulomics nanobind tissue kit (102-302-100) with some modifications to account for tighter packing of the sperm genome. We supplemented the lysis solution with 150mM 1,4-Dithiothreitol (DTT) to disrupt protamine disulfide bonds and extended proteinase K incubation from 30 minutes to 2 hours. The rest of the Circulomics HMW DNA extraction protocol remains unchanged. HMW DNA was sheared to 10-14 kb DNA fragments using Megaruptor 3 system (B06010003) with speed setting 30. CCS sequencing libraries were constructed according to the standard CCS library preparation protocol 1.0 (100-222-300), and the libraries were sequenced using a Sequel IIe instrument at the Wellcome Sanger Institute.

For the SL data, samples were thawed on ice and cells were pelleted in an isotonic sperm wash solution to remove debris and reduce somatic cell contamination. DNA was extracted using the NEB Monarch® HMW DNA Extraction Kit for Cells & Blood (#T3050) UHMW protocol, with some modifications to the cell lysis step. Specifically, cells were digested at 56°C for 1 hour at 300 rpm with 100 µL Nuclei Prep Buffer, 100 µL Nuclei Lysis Buffer, 10 µL Proteinase K (NEB #P8107, 20 mg/ml), and 10 µL 1M DTT (GoldBio #DTT in dH2O, final concentration ∼50 mM). An additional 20-minute digestion was then performed with 5 µL RNase A (NEB #T3018, 20 mg/ml). The rest of the protocol remained unchanged. HudsonAlpha performed HMW DNA quality control, CCS library preparation, and sequencing on the PacBio Revio sequencing instrument.

### De-novo haplotype-resolved assembly

We used `hifiasm`^28^ (version 0.19.5-r592, default parameters, HiFi only mode) to construct a haplotype-resolved de-novo assembly for each sample. The `hifiasm` assembler outputs a partially phased assembly graph for the two haplotypes, which we converted into two fasta sequence files per sample.

We scaffolded each haplotype onto the T2T-CHM13 reference genome^39^ with RagTag^40^ (version v2.1.0, with arguments “`-u -w --aligner minimap2`”). RagTag orients and places contigs along the T2T reference with gaps; we manually expanded the gaps between adjacent contigs to 30000N to avoid reads mapping across a gap to two contigs, as this may include a phase switch error which will further be falsely interpreted as a CO.

### PRDM9 genotyping

We obtained the zinc-finger array sequences from 74 PRDM9 alleles reported previously^25^ and mapped them to each assembled haplotype of each donor. For each haplotype, we called the PRDM9 allele as the one with the smallest amount of mismatches, insertions or deletions between the allele sequence and the assembly. All the A-alleles, B-allele and the D-allele were a perfect match, while we did not find a perfect match to the new PRDM9 allele of AN-s1/2. In addition, we verified that the flanking regions on the two sides of the sequence match the expected sequences as reported^25^ (up to 2bp mismatches), and confirmed that PRDM9 allele variations are not due to assembly errors by visually inspecting the assembly around the region.

### Read alignment

CCS reads with residual adapter sequences were identified with HiFiAdapterFilt^41^ and were excluded from downstream sequence analysis. We used `minimap2`^42, 43^ (version 2.26-r1175, with arguments “`-ax map-hifi --cs=short --eqx --MD`”) to map the reads to each haplotype separately. We then filtered each bam file to contain only primary alignments (using the “`-F 0x900`” flag in “`samtools` `view`”^44^) and with mapping quality MAPQ=60. We further discarded reads that map to only one of the haplotypes, or that map to different chromosomes. Finally, we discarded reads that map to different strands of the two haplotypes.

### Creating candidate recombinant reads

#### Comparing haplotype alignments

Each alignment of a read to a haplotype effectively partitions the read sequence into segments, where each segment corresponds to an alignment operation - either (i) “match” - aligns and matches perfectly to the reference haplotype; (ii) “mismatch” - aligns but does not match to the reference; (iii) an “insertion” (iv) a “deletion” with respect to the reference (in a deletion, this read segment is of length 0); or (v) a “soft clipping”.

We used the two alignments of the same read to the two haplotypes, and created a refined partition of the read according to both alignments, defined as the partition which is the intersection of the two partitions. In this refined partitioning, each read segment corresponds to two alignment operations. For example, the read segment [0, 40) is both “match” to haplotype 1 and “match” to haplotype 2; the read segment [40, 41) is a “match” to haplotype 1 but “mismatch” to haplotype 2; the read segment [41, 41) is a “deletion” with respect to haplotype 1, but is an “match” (in an empty sense) to haplotype 2; the read segment [41, 50) is a “match” to haplotype 1 but an “insertion” with respect to haplotype 2, and so forth. This refined joint partition is the basis for our followup analysis.

We discarded reads with more than 10bp of soft clipping to either haplotype, as these were observed to often be chimeric reads. Moreover, we discarded reads with more than 100 sequence errors, defined as segments (usually 1bp) mismatching both haplotypes.

#### SNP calling and filtering

Along each read, we find SNP candidates using the partitioning described above to find single nucleotide positions which match to one haplotype but do not match to the other haplotype. We observed that many such one nucleotide mismatches occur in regions suspicious of assembly errors, such as regions of low complexity, tandem repeats, near read ends and in haplotype regions with low coverage. Therefore, we further filtered SNP candidates according to several criteria.

First, we include only those surrounded by at least 10bp of matched alignment on both sides of the SNP. Second, we discarded SNPs that were within 1500bp of either read ends for the Sequel II data; or 400bp for the Revio data (see Supplementary Note about read trimming calibration). Third, we filtered SNPs with a base quality (BQ) score of less than 60 for the Sequel II data; or 40 for the Revio data (see Supplementary Note about BQ threshold calibration). Fourth, we ran `sdust`^45^ (version 0.1-r2, with default arguments), an implementation of the dustmasker algorithm^46^, to identify low complexity regions in both haplotypes. We filtered SNPs that were mapped to a low complexity region in either haplotype. Fifth, we ran Tandem Repeat Finder^47^ (version v4.09.1, with arguments “`2` `6` `6` `80` `10` `50` `500` `-ngs -h -l 10`”) to find tandem repeats in both haplotypes. We filtered SNPs that mapped to a tandem repeat in either haplotype. SNPs that passed these criteria are considered high confidence SNPs. We discarded reads that have no such high confidence SNPs from further analysis.

For the Sequel II data, we called a further set of candidate SNPs satisfying the same criteria, except for a minimal BQ of 30 (instead of 60 in high confidence SNPs) and a distance of more than 500bp from read ends (vs. 1,500bp in high confidence SNPs) as a less strict subset of “classification SNPs’’ we used for event classification and analysis, but not for event detection. For the Revio data, we used a minimal BQ of 30 (vs. 40) and distance of 200bp (vs. 400bp).

#### Haplotype assignment

We scanned each read for high confidence SNPs and kept track of which of the two haplotypes each SNP matches to. The large majority of reads (>99.9%) have SNPs that are fully consistent with at least one of the haplotypes, and therefore are not candidates to be recombinant reads. However, their existence is helpful in filtering assembly errors, as we describe below. Therefore, we recorded, for each of the two haplotypes and for each read, the percentage of SNPs in the read consistent with the haplotype.

#### Further SNP filtering measures

We observed that regions in an assembled haplotype that did not have enough consistent reads overlapping them were prone to errors and false detection. We therefore further require that each SNP have at least three reads overlapping the focal SNP, which have >95% SNPs consistent with haplotype 1; and similarly for haplotype 2.

#### Recombinant SNP candidates

To create a list of candidate recombinant reads, we calculated again for every read the percentage of high-quality SNPs (after read coverage filtering) consistent with each one of the haplotypes. A read is therefore a candidate if its SNPs are not fully consistent with either haplotype.

### Classifying candidate recombinant reads

#### Further read filtering

Based on the list of classified reads and their annotations, we apply further filtering to reduce false positives. We define a transition pair as a pair of adjacent SNPs on the read which map to different haplotypes. Each candidate read will have at least one transition pair. Two transition pairs (across different reads) that map to the same haplotype coordinates are suspected of being a result of a phasing or assembly error, since we assume every recombination event happens exactly in one read. We therefore discarded reads if they include transition pairs recurring in other reads.

In addition, if the genomic region between a transition pair is not covered by three or more reads (see “Further SNP filtering measures” above) in either haplotype, then we deem the assembly in this region as uncertain. In particular, such low coverage regions may indicate a phasing error in haplotype assembly, which may result in false CO calls. We therefore also discard reads with such low coverage transition pairs.

#### Event classification

We classify each candidate read as CO is there is only one transition pair, flanked by at least one SNP on either side (e.g. “1122” - that is, two SNPs from haplotype 1 followed by two SNPs from haplotype 2); as NCO is there are two transitions (e.g. “121”); as “ambiguous”, indicating we cannot distinguish CO and NCO, if there is one transition but at least one side without flanking SNPs (e.g. “12”, “112”, or “122”); or “complex” is there are more than two switches.

#### Mapping to reference genomes

For genetic distances, we used the European male-specific refined genetic map^30^, which is provided in GRCh37 coordinates. To analyse CO rate information, we mapped all reads to the GRCh37 reference genome^48^ using `minimap2`^42, 43^ (version 2.26-r1175, with arguments “`-ax map-hifi --cs=short --eqx --MD`”). For DSB hotspots, the DSB map^31, 32^ is provided in GRCh38 coordinates. To analyse DSB data, we similarly used `minimap2` to map all reads to the GRCh38 reference genome^49^. To analyse the distance to telomeres, we also mapped all reads to the T2T reference genome^39^.

#### H3K4me3 map

We downloaded histone ChIP-Seq peak data (ENCODE3 GRCh38) obtained from an adult male testis tissue (37 years), from the ENCODE portal^36, 50, 51^ (https://www.encodeproject.org/) with the identifiers ENCSR611DJQ and ENCFF537BJO.

#### Testing for equality of distributions

We first describe how we test for equality of distribution between two samples. Consider two sets of continuous measurements (e.g. recombination rates) for two samples. We use the Anderson-Darling (AD) statistic^52^ for two-sample equality of distribution testing, which is based on the integral of the square differences between the empirical cumulative distribution functions of the two measurement sets. To calculate the statistical significance of the observed AD statistic, we perform permutation testing by randomly assigning each measurement to one of the samples, while retaining sample sizes. We permute the data 10,000 times, calculate the AD statistic each for each permutation, and record the fraction of permutations for which the AD statistic was larger than the observed AD statistic as our P-value.

To test for equality of distribution between a set of measurements from one sample, and several sets of measurements from other samples, we first calculate the AD statistic between the focal sample and each other sample, and take their sum as our statistic. To calculate statistical significance, for 10,000 times we perform the permuted assignment as above separately to the focal sample and each of the other samples, and calculate the sum of AD statistics across the randomised sets. Our P-value is the fraction of (pairwise sequences of) permutations resulting in a statistic larger than the observed one.

### Software

We used snakemake^53^ v7.32.4 for pipeline management and execution and polars v0.20.13 for large-scale data analysis.

## Data availability statement

All data relating to TwinsUK samples have been deposited to the TwinsUK BioResource data management team and are available by application to the Twin Research Executive Access committee (TREC) at King’s College London. The TwinsUK BioResource is managed by TREC, which provides governance of access to TwinsUK data and samples. TwinsUK data users are bound by data sharing agreement set out in the data access application form, which includes responsibilities with respect to third party data sharing and maintaining participant privacy. Further responsibilities include a responsibility to acknowledge data sharing. EGA submission of sequence data is in progress.

## Code availability statement

The code used to generate the data and analysis will be available at publication.

## Acknowledgements

We thank the participants and local coordinators at the TwinsUK, the staff of Wellcome Sanger Institute long-read sequencing team for CCS library preparation and sequencing, the ENCODE Consortium for the H3K4me4 data, and Anders Charmouh, Mikkel Schierup and Zachary Baker for constructive discussions and information and Anjali Gupta Hinch for communication about the double-stranded break data we used. For the purpose of Open Access, the authors have applied a CC BY public copyright licence to any Author Accepted Manuscript version arising from this submission.

## Authors contributions

R.D. and R.R. designed the experiments. K.S. and S. Lee, and S. Li modified protocols and performed HMW DNA extraction from sperm samples. R.S. and S.Lee performed data quality control and initial analysis. R.S. carried out further analysis including statistical inference and

C.Z. carried out the genome assemblies. T-P.Y. helped with PRDM9 genotyping. A.N., S.W. and K.S.S. designed and carried out sample recruitment and collection. S.Li and P.H.S. provided additional sequence data. E.M. and P.J.C. provided blood samples. R.S., S.Lee and R.D. wrote the manuscript. All authors reviewed and edited the manuscript.

## Extended Data Figures

**Extended Data Figure 1.**
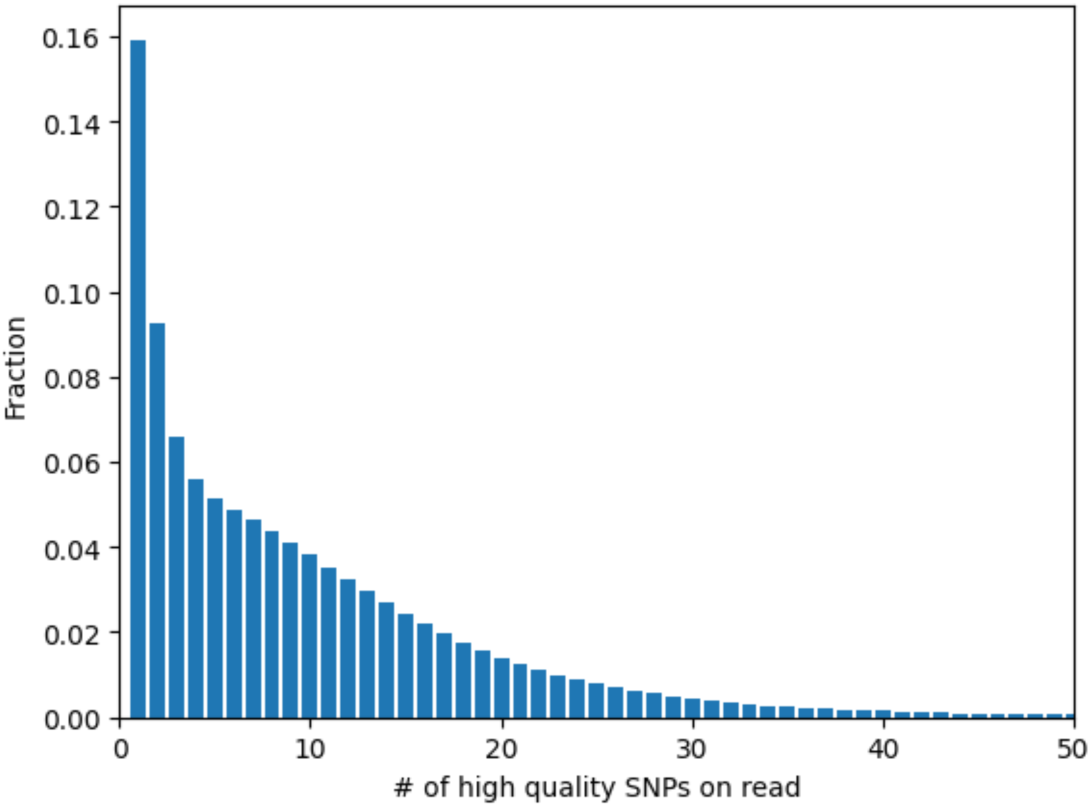
The distribution of the number of SNVs passing quality control on reads.

**Extended Data Figure 2.**
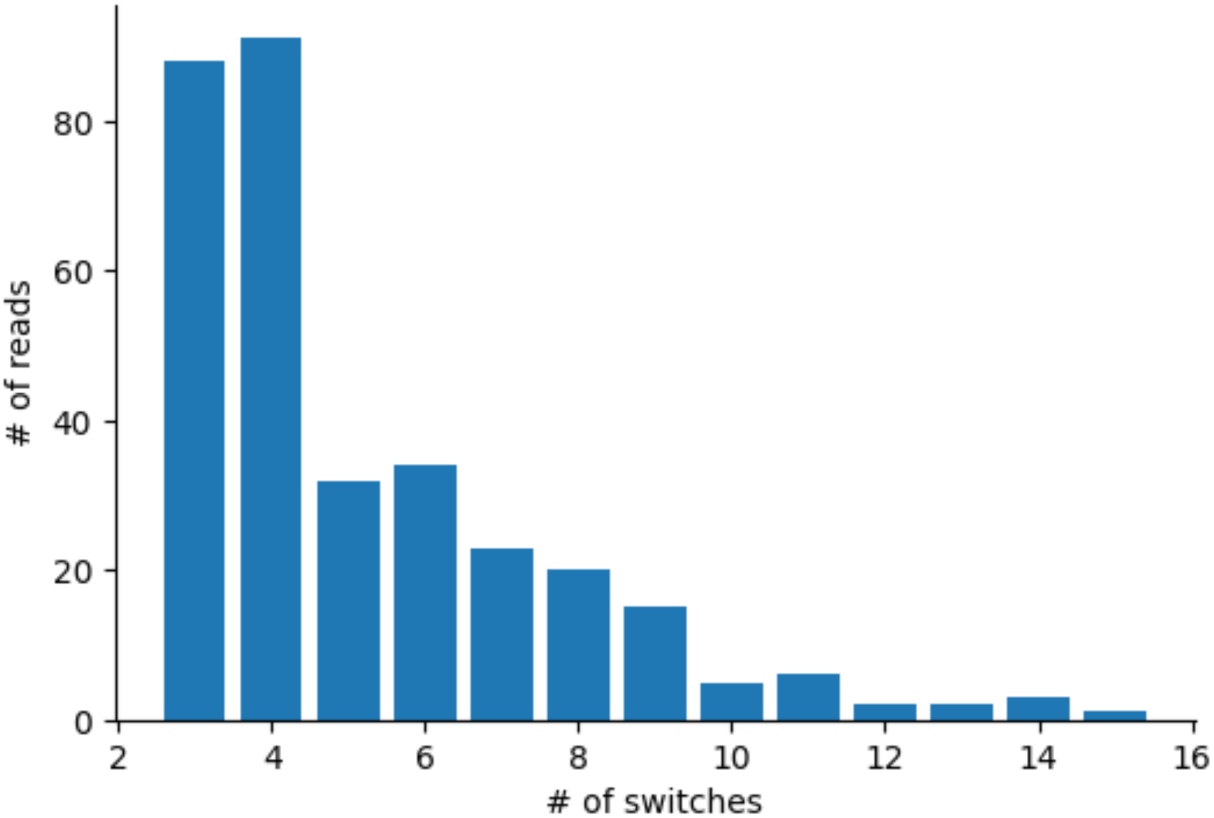
The distribution of the number of observed switches in reads with complex recombination events.

**Extended Data Figure 3.**
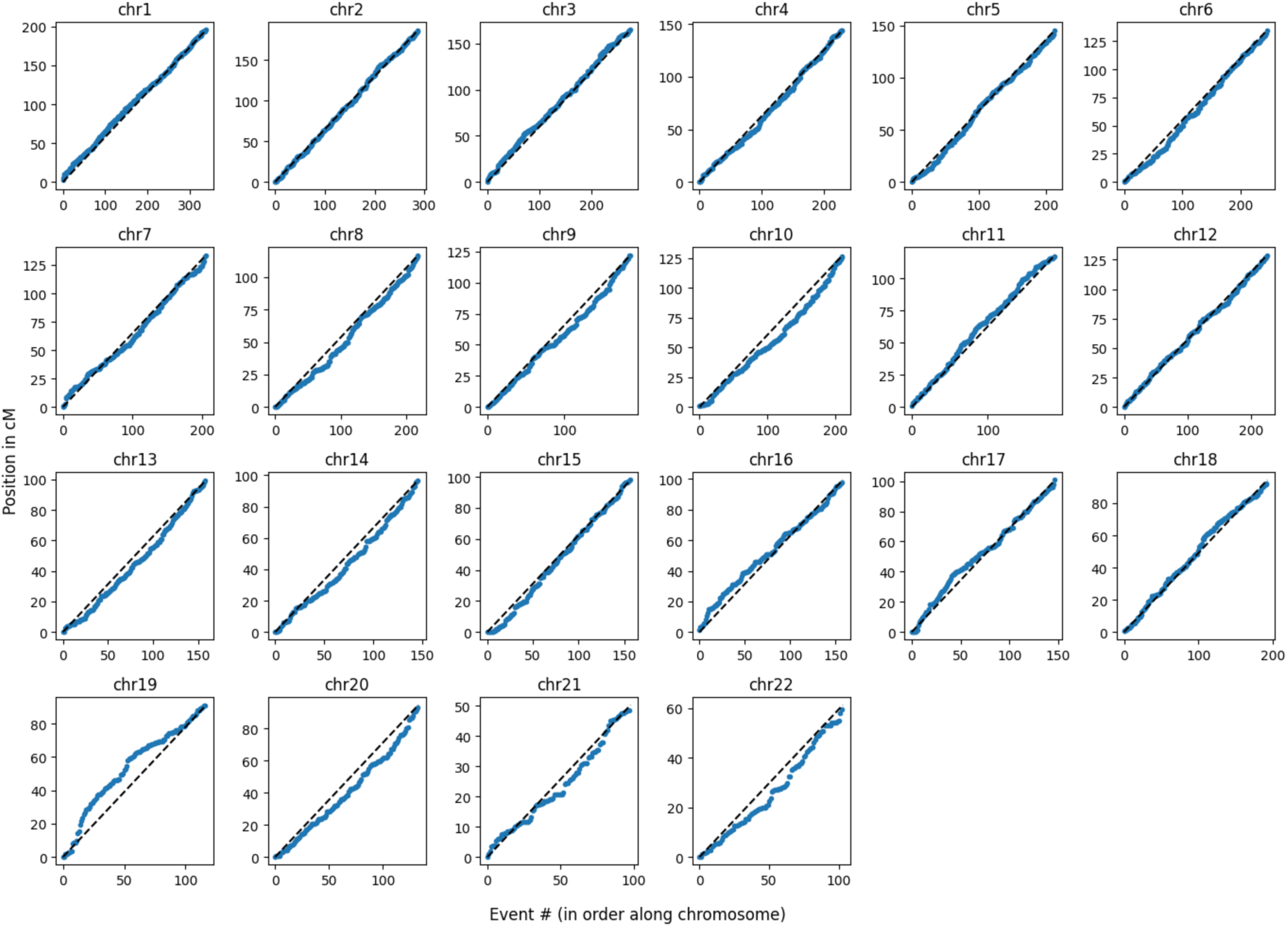
The positions of CO reads along the chromosomes, in genetic coordinates.

**Extended Data Figure 4.**
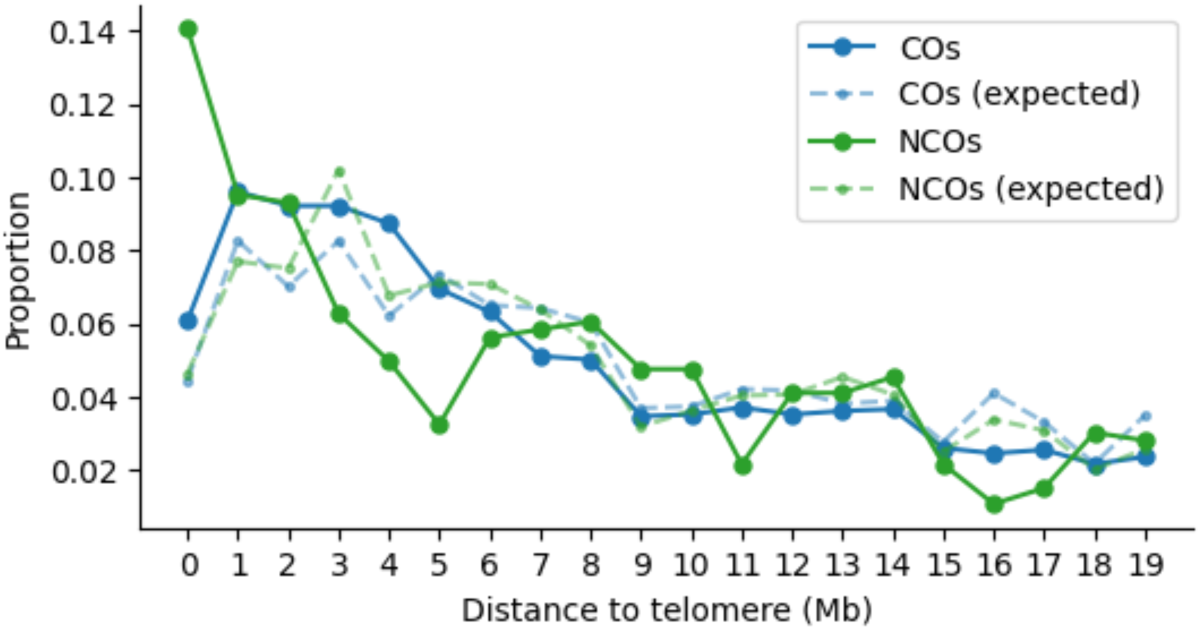
The distribution of distances of CO and NCO events from telomeres, for events that occur within 20Mb of chromosome ends, and excluding acrocentric chromosomes. The expectation is calculated taking into account the CO recombination rates and the marker patterns on reads, to account for hotspots and marker density.

**Extended Data Figure 5.**
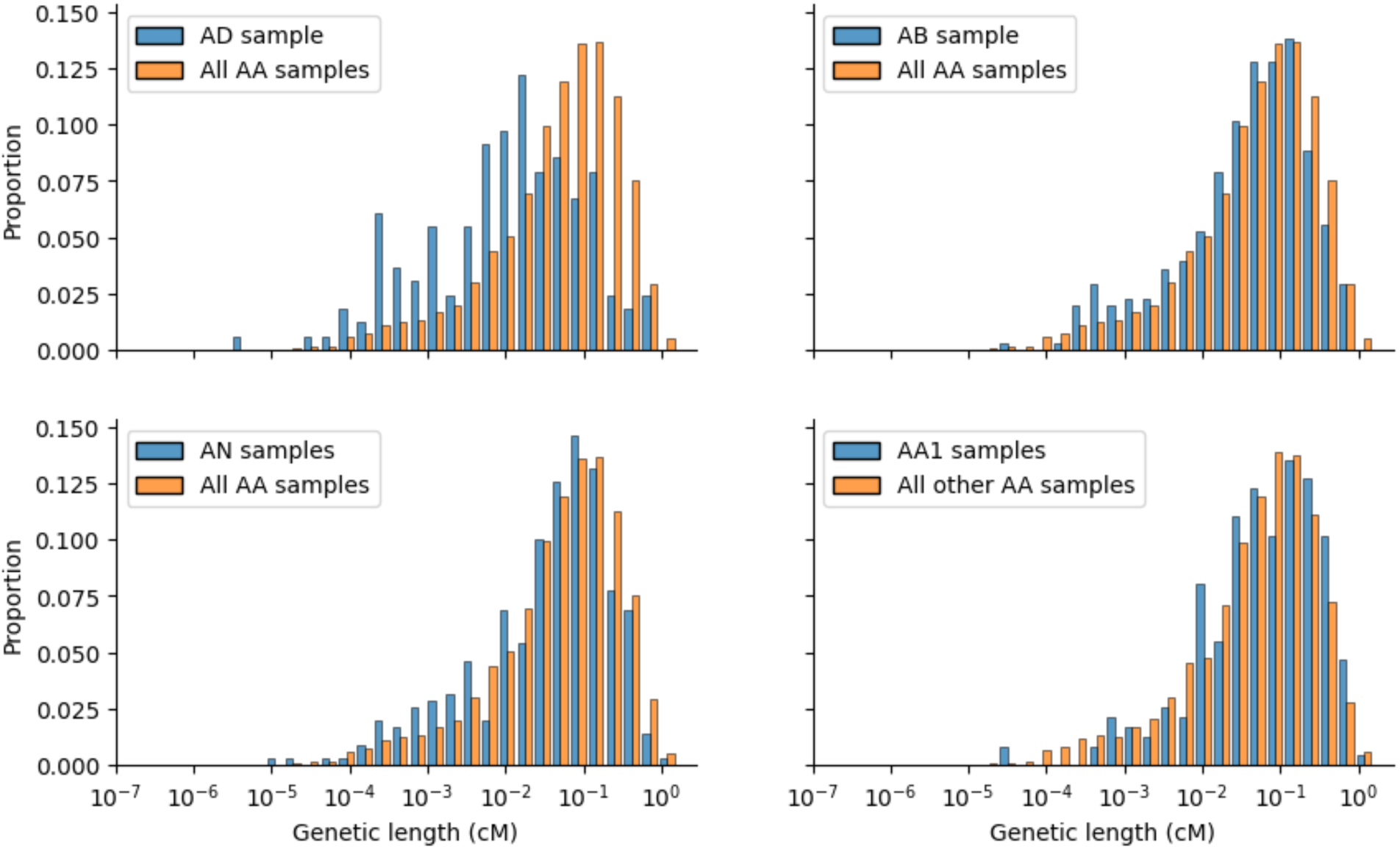
The difference in distribution of genetic length (in cM) of CO reads between: (A) the AD sample and the A/A samples; (B) the AB sample and the A/A samples; (C) the AN samples and the A/A samples; (D) the AA1 samples and the other A/A samples.

**Extended Data Figure 6.**
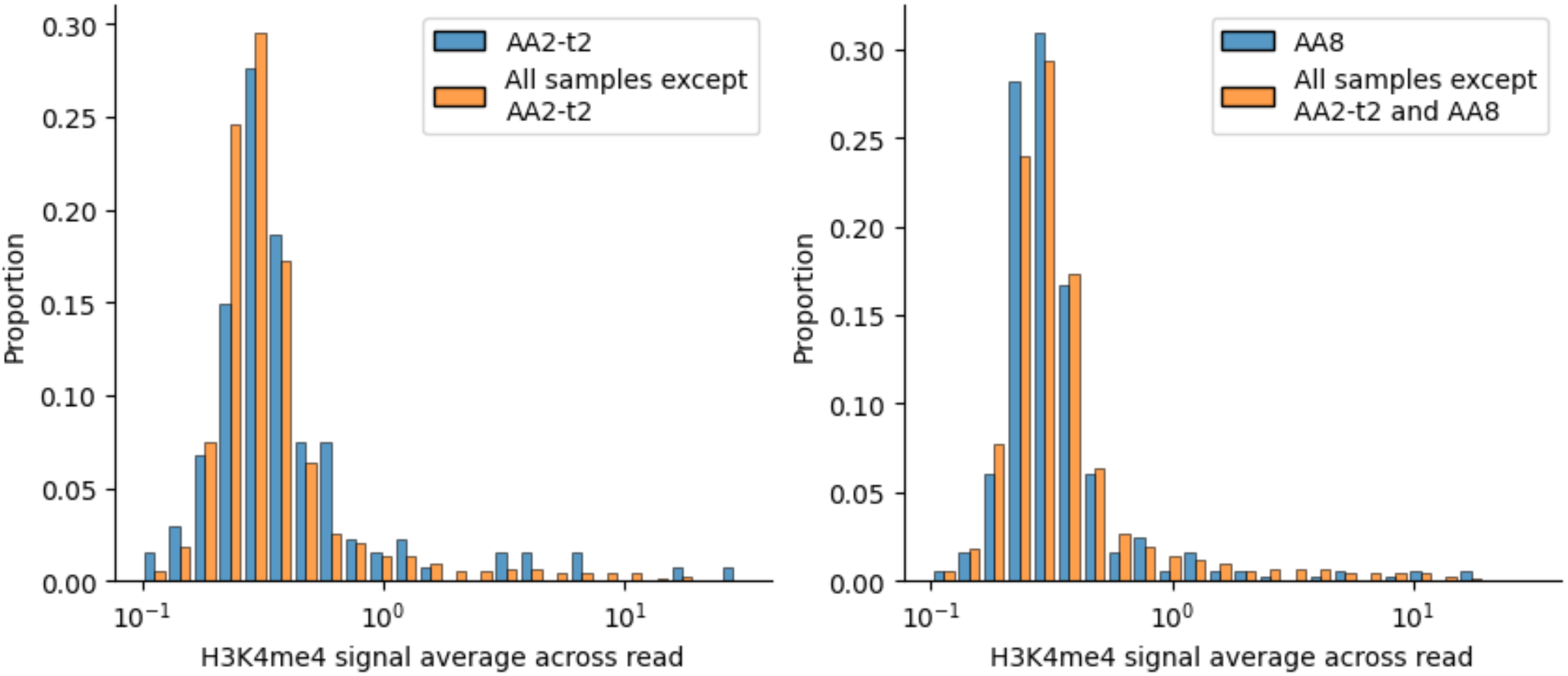
The difference in distribution of H3K4me3 signal average across CO reads between (A) the AA2-t2 sample and all the other samples; (B) the AA8 sample and all other samples except AA2-t2.

**Extended Data Figure 7.**
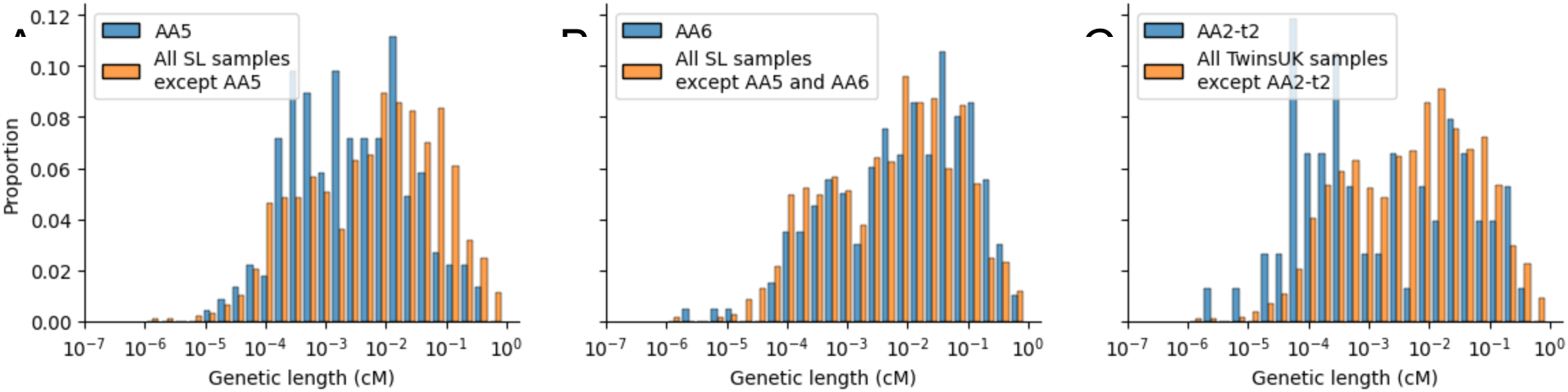
The difference in distribution of genetic length (in cM) of NCO reads between (A) the AA5 sample and all other SL samples; (B) the AA6 sample and all other SL samples except AA5; (C) the AA2-t2 sample and all other TwinsUK samples.

**Extended Data Figure 8.**
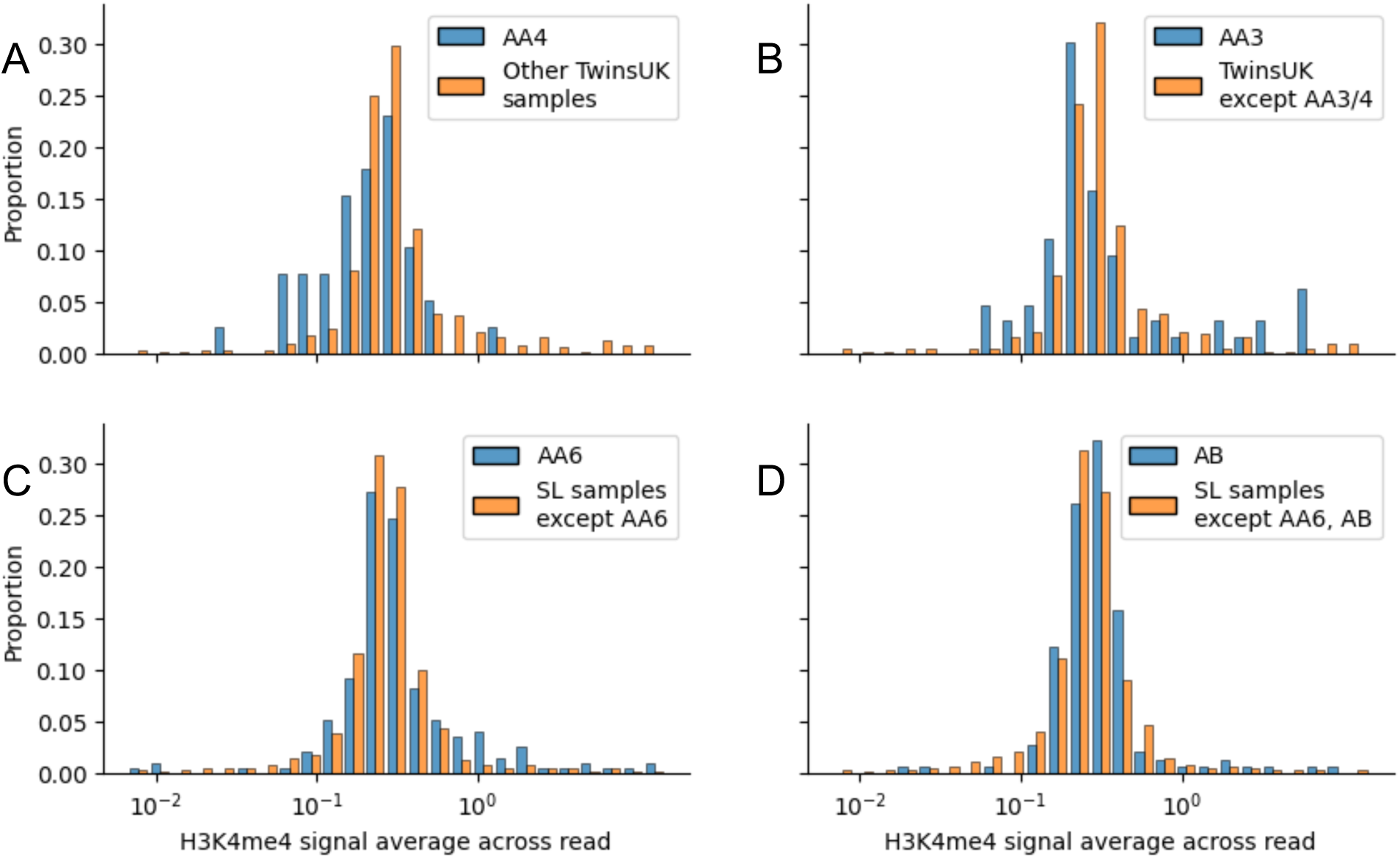
The difference in distribution of H3K4me3 signal average across NCO reads between (A) the AA4 sample and all other TwinsUK samples; (B) AA3 and other TwinsUK samples except AA4; (C) AA6 and other SL samples; (D) AB and other SL samples except AA6.

