## Supplementary Information for "Insights into non-crossover recombination from long-read sperm sequencing"

### Supplementary Information for
#### *Schweiger et al. "Insights into non-crossover recombination from long-read sperm sequencing"*

#### Calibration of read trimming

For the subset of reads of minimal length 5kbp, we counted the number of 1bp segments which are mismatches to both haplotypes, as a function of the distance to read end (<5kbp, bins of 10bp). Since some of these mismatches are a result of assembly errors, we further limited our analysis to high confidence 1bp mismatches with; at least 10bp of matched alignment on both sides of the segment; do not overlap a low complexity or a repeat (Methods); and with a BQ of 30 or more (we need only choose a reasonable BQ level, since for this analysis we only care about relative error rates between read end distances).

For the TwinsUK samples, sequenced by the Sequel II instrument, we observed a slow decay of error rates until a distance of 2-3kbp from read end (Supplementary Fig. 1). We selected a threshold of 1,500bp for detection SNPs (that is, SNPs within 1,500bp of read ends would not be used to detect recombination events), and a less stringent threshold of 500bp for classification SNPs (used to classify events, once detected).

For the SL samples, sequenced by the Revio instrument, we observed a substantially lower baseline error rate, as well as a faster drop from read ends to the baseline (Supplementary Fig. 2). We therefore used a threshold of 400bp for detection SNPs and 200 for classification SNPs.

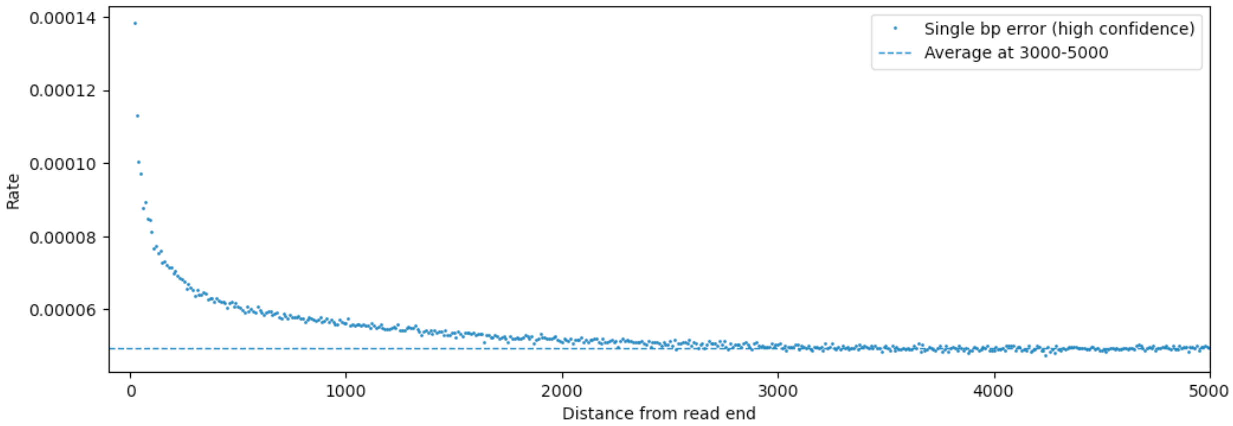


**Supplementary Figure 1.** Relative rates of high confidence single-bp mismatches, as a function of distance to read ends, for the Sequel II instrument.

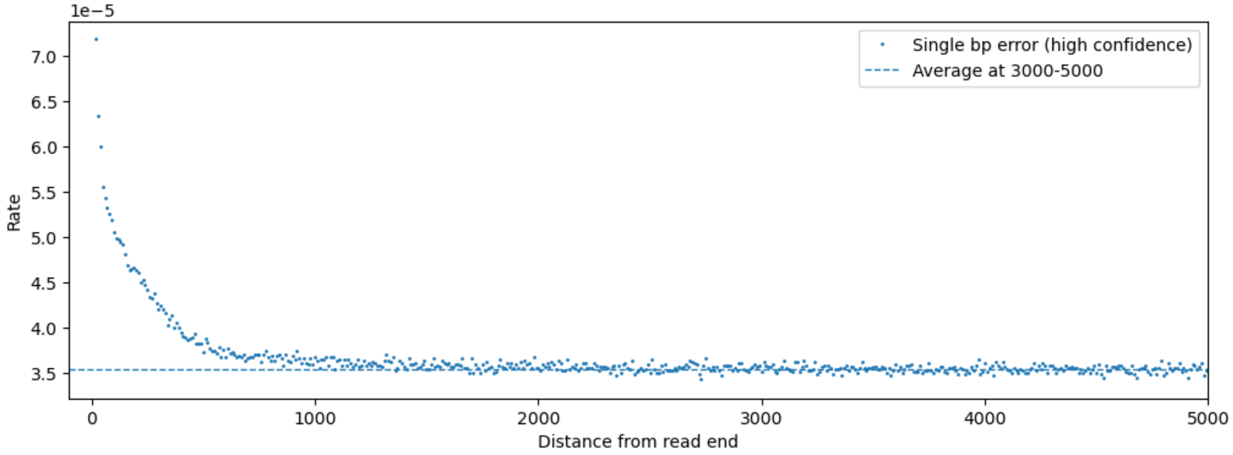


**Supplementary Figure 2.** Relative rates of high confidence single-bp mismatches, as a function of distance to read ends, for the Revio instrument.

#### Calibration of base quality scores

To calibrate the base quality (BQ) scores, we make use of the high coverage of reads to estimate whether a mismatch to a haplotype is a gene conversion or a sequencing error, and therefore get an estimate of false positive NCO calls. In more detail, we analyse SNPs - 1bp segments that match one haplotype but mismatch the other; that are not within 1,500bp of the read end; have at least 10bp of matched alignment on both sides of the segment; do not overlap a low complexity region or a repeat (see Methods). We further limit our analysis to reads where the 1bp segment mismatch is surrounded by at least 3 SNPs which do match the haplotype (i.e. a pattern of "...1112111...").

These 1bp SNPs can be caused either by NCO leading to gene conversion; by a sequencing error, or by de-novo mutations. Of these, the latter is rare (a rate of $10^{-8}$/bp), so these SNPs are likely to originate from either an NCO or a sequencing error. To distinguish between these two scenarios, we compare the SNP to the other haplotype. If this is a sequencing error, there is no particular reason that it should flip to the other allele vs. any other nucleotide, apart from the particular sequencing error patterns. Conversely, if this is a true GC, then it should always flip to the other allele. Therefore, the ratio between the number of SNPs that flip to a nucleotide different than the two haplotypes, and the number of SNPs that flip to the allele of the other haplotype, is a proxy of the false positive call rate we expect for NCOs.

For Sequel II data (Supplementary Fig. 3), we set the BQ threshold for detection SNPs to be 60, and 40 for classification SNPs. For Revio data (Supplementary Fig. 4), the BQ threshold for detection SNPs was 40 and 30 for classification SNPs.

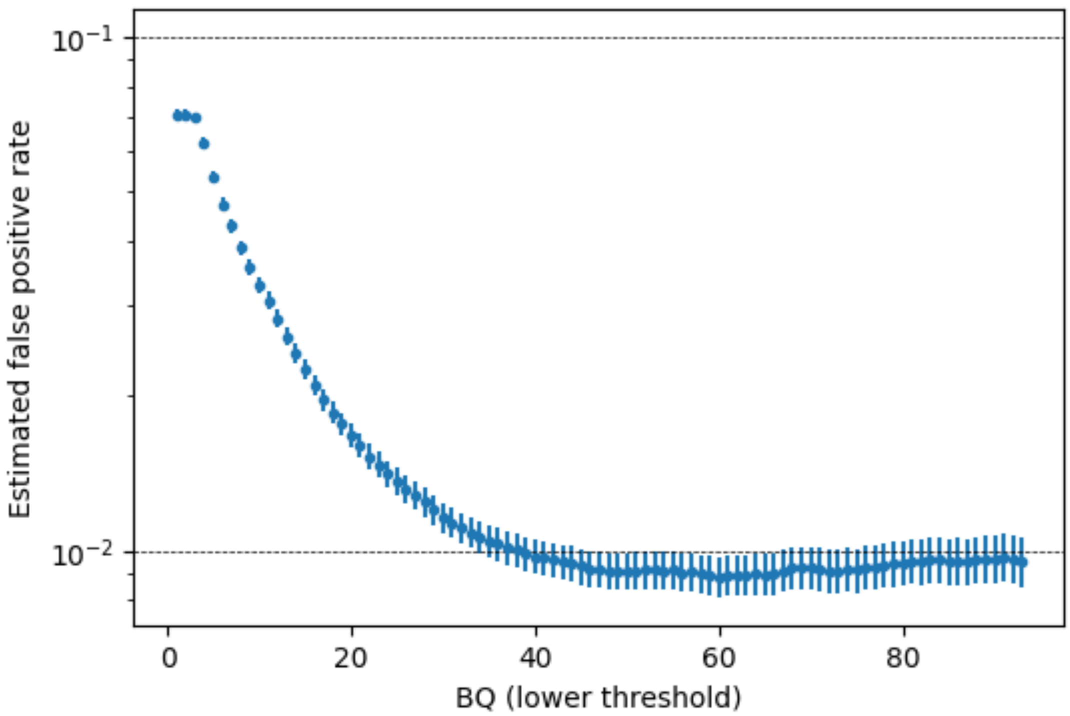


**Supplementary Figure 3.** An estimate of the rate of false positive NCO calls, as a function of base quality (BQ) minimal threshold, for Sequel II data.

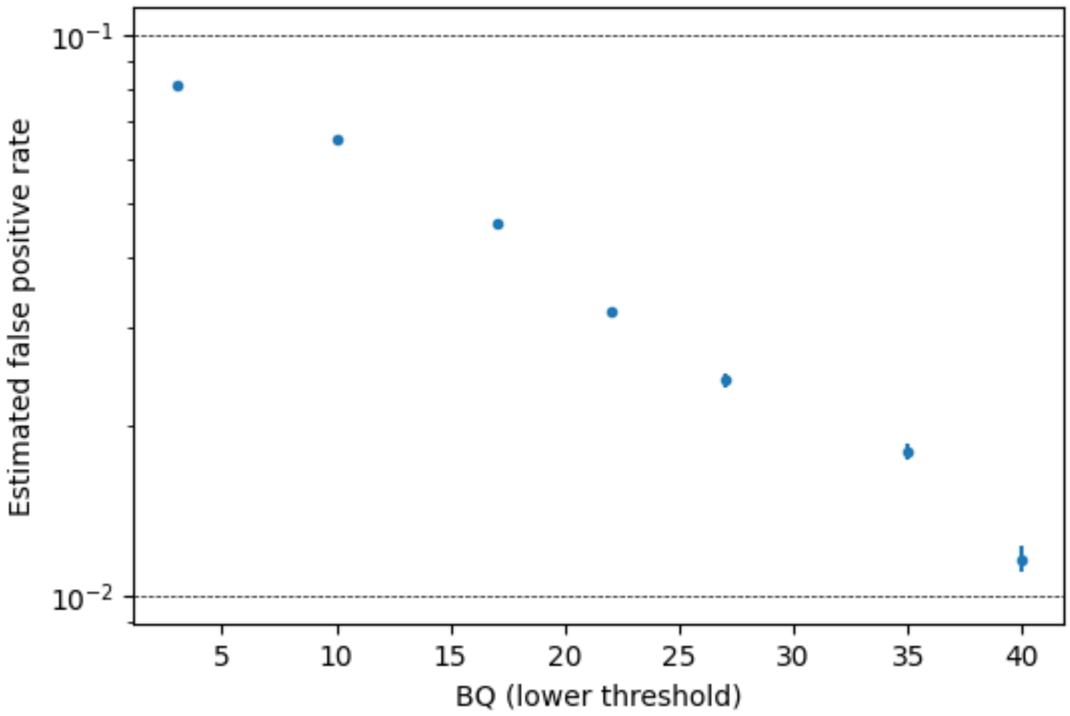


**Supplementary Figure 4.** An estimate of the rate of false positive NCO calls, as a function of base quality (BQ) minimal threshold, for Sequel II data.

### Testing for differences in CO and NCO event distributions

#### Problem setup

We would like to test if the spatial distribution of CO rates along the genome is different from that of NCO events. The direct way to do this would be to observe a large number of both CO and NCO events everywhere along the genome and compare them at each location. However, this would require unrealistically high coverage, many thousands if not millions of times higher than we have. Therefore, instead we group together reads that are expected to behave similarly in terms of recombination and detection probabilities, and then test for differences between groups of CO reads and NCO reads, as we describe below. The CO recombination map is a natural proxy for such grouping, as CO rates are expected to follow it closely.

We first limit ourselves to reads (in addition to QC described above) with at least 5 SNPs, and we define the genomic interval between the second SNP and the second-to-last SNP as the *detectable interval*. We will only be interested in CO reads where the recombination occurs in the detectable interval, i.e. the first two (or more) SNPs belong to one haplotype and the last two (or more) SNPs to other ("11...22"). Similarly, we will only be interested in NCO reads where the converted SNP/s are in the detectable interval, i.e. the first and last two SNPs are not converted ("11...2...11"). This increases the confidence of correct classification while dealing with the same set of SNPs for both types of events.

Our null model assumes that (a) both CO and NCO events are initiated by a DSB, which is initiated according to some spatial distribution along the genome; (b) each DSB site is resolved to a CO at probability $q$ and to an NCO at probability $1-q$, independent and uniformly along the genome; (c) COs occurring in the detectable interval are always detected, while NCOs will only be detected if a marker within the detectable interval is converted; (d) NCO tract lengths are geometrically distributed with a known mean length, and therefore, given the pattern of markers on a read, we can calculate the probability of detection. Note that for this analysis we assume a single geometric distribution for tract lengths and not a mixture, for simplicity.

Formally, we use the following graphical model (illustrated in Supplementary Fig. S5) as our null model:

1. The genetic length $L_i$ of a detectable interval is sampled from a random variable $L$.
2. A recombination event occurs in the detectable interval ($R_i=1$) according to a probability $u(L_i)$ determined by the genetic length $L_i$.
3. The type $T_i \in \{CO, NCO, None\}$ of the recombination event is determined independently for each read conditional on $R_i$; if $R_i=1$ then it resolves to a CO with probability $q$, and to a NCO with probability $1-q$; and if $R_i=0$ it is always $None$. The SNP pattern $P_i$, which consists of the number of SNPs and their distances along the detectable interval, is sampled from a random variable P, which in general is not independent of L.

4. The random variable $D \in \{CO, NCO, None\}$ determines if and which type of recombination event was detected. If no event occurred ($R_i = 0$ and $T_i = None$), then also $D_i = None$. If a CO occurred ($T_i = CO$) at the detectable interval, then it is always detected and $D_i = CO$. If a NCO occurs ($T_i = NCO$), then it is detected in a probability $f_i$ determined only by the SNP pattern $f_i = f(P_i)$.
5. If the NCO tract lengths are geometrically distributed with a known mean tract length, we can calculate $f_i$ explicitly according to the likelihood model described in the main text.
6. *L, P* and *D* are observed, the rest of the variables are not.

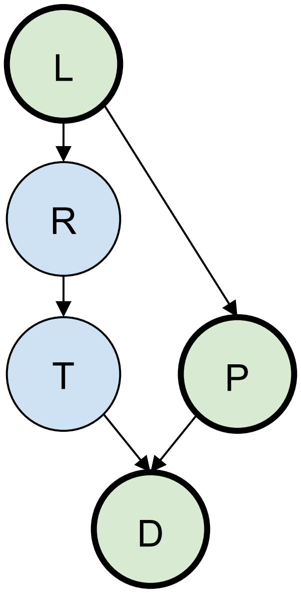


**Supplementary Figure 5.** The graphical model underlying the test for differences between COs and NCOs. The genetic length (*L*) is sampled from a random variable; a recombination event occurs (*R*) in probability determined by *L*; the type (*T*) of recombination event is determined; then this event is detected (*D*) in probability determined by its type *T* and the SNP patterns on the read (*P*).

In terms of this graphical model, the assumption that the decision to resolve into CO or NCO is independent of the genetic coordinates translates into $T \perp L \mid R$ (i.e. *T* is independent of *L*, conditional on *R*; indicated in the diagram as T just depending on R). However, we do not observe *T* directly, only via *D*, which is affected by *L* via *P*.

#### Test: Assuming independence between cM rates and SNP patterns

The simplest assumption to make is that the probability of recombination in a candidate interval $u(l)$ is independent of the chance of an NCO being detected in that interval if it happened, $\sum_p f(p)\, Pr\,(p)$. In terms of the graphical model, we assume here $P \perp L$. In this case, we can decompose:

$$Pr(D = CO \mid L = l) = \sum_{r,t} Pr(D = CO \mid T = t) \cdot Pr(T = t \mid R = r) \cdot Pr(R = r \mid L = l)$$

Since a CO will only be detected if is occurred, this is:

$$= Pr(D = CO|T = CO) \cdot Pr(T = CO \mid R = 1) \cdot Pr(R = 1|L = l) = 1 \cdot q \cdot u(l)$$

Similarly for NCOs,

$$Pr(D = NCO \mid L = l) = \sum_{r,t,p} Pr(D = NCO \mid t,p) \cdot Pr\,(p) \cdot Pr(\,t \mid r) \cdot Pr(r \mid l)$$

$$= (1 - q)u(l) \cdot \sum_{p} f(p)\, Pr\,(p)$$

Using Bayes' law,

$$Pr(L = l|D = CO) = Pr(D = CO \mid L = l) \cdot Pr(L = l)/Pr(D = CO) \propto u(l)Pr(L = l)$$
$$Pr(L = l|D = NCO) = Pr(D = NCO \mid L = l) \cdot Pr(L = l)/Pr(D = NCO) \propto u(l)Pr(L = l)$$

It follows that these two conditional distributions are identical, and can therefore be tested with a two-sample test for equality of 1D distributions, e.g. Kolmogorov-Smirnov or Anderson-Darling.

Because we observe L and P directly, and f is a function of P, we can examine whether the assumption of L and f being independent is reasonable. They are indeed significantly correlated, but very weakly (Pearson's r=0.02, p<1e-100). Furthermore, the correlation is positive, which would make NCOs more detectable for higher values of L, whereas we see in Fig. 2 that NCOs are relatively less detected at higher values of L. The slight positive correlation may be due to the mutagenic effect of recombination which may lead to a positive correlation between cM rates and SNP density.

### Sequence of the new PRDM9 allele

TGTGGACAAGGTTTCAGTGTTAAATCAGATGTTATTACACACCAAAGGACACATACAGGGGAGAAGCTCT
ACGTCTGCAGGGAGTGTGGGCGGGGCTTTAGCTGGAAGTCACACCTCCTCATTCACCAGAGGATACACAC
AGGGGAGAAGCCCTATGTCTGCAGGGAGTGTGGGCGGGGCTTTAGCTGGCAGTCAGTCCTCCTCACTCAC
CAGAGGACACACACAGGGGAGAAGCCCTATGTCTGCAGGGAGTGTGGGCGGGGCTTTAGCTGGCAGTCAG
TCCTCCTCACTCACCAGAGGACACACACAGGGGAGAAGCCCTATGTCTGCAGGGAGTGTGGGCGGGGCTT
TAGCCGGCAGTCAGTCCTCCTCACTCACCAGAGGAGACACACAGGGGAGAAGCCCTATGTCTGCAGGGAG
TGTGGGCGGGGCTTTAGCCGGCAGTCAGTCCTCCTCACTCACCAGAGGAGACACACAGGGGAGAAGCCCT
ATGTCTGCAGGGAGTGTGGGCGGGGCTTTAGCTGGCAGTCAGTCCTCCTCAGTCACCAGAGGACACACAC
AGGGGAGAAGCCCTATGTCTGCAGGGAGTGTGGGCGGGGCTTTAGCTGGCAGTCAGTCCTCCTCACTCAC
CAGAGGACACACACAGGGGAGAAGCCCTATGTCTGCAGGGAGTGTGGGCGGGGCTTTAGCAATAAGTCAC
ACCTCCTCAGACACCAGAGGACACACACAGGGGAGAAGCCCTATGTCTGCAGGGAGTGTGGGCGGGGCTT
TCGCGATAAGTCACACCTCCTCAGACACCAGAGGACACACACAGGGGAGAAGCCCTATGTCTGCAGGGAG
TGTGGGCGGGGCTTTAGAGATAAGTCAAACCTCCTCAGTCACCAGAGGACACACACAGGGGAGAAGCCCT
ATGTCTGCAGGGAGTGTGGGCGGGGCTTTAGCAATAAGTCACACCTCCTCAGACACCAGAGGACACACAC
AGGGGAGAAGCCCTATGTCTGCAGGGAGTGTGGGCGGGGCTTTCGCAATAAGTCACACCTCCTCAGACAC
CAGAGGACACACACAGGGGAGAAGCCCTACGTCTGCAGGGAGTGTGGGCGGGGCTTTAGCGATAGGTCAA
GCCTCTGCTATCACCAGAGGACACACACAGGGGAGAAGCCCTACGTCTGCAGGGAG

### Supplementary Tables

| Sample ID | PRDM9 genotype | Dataset | Age | # of CCS reads | Read length ± std | Read depth |
|---|---|---|---|---|---|---|
| AA1-s1 | A/A | TwinsUK | 62 | 3,947,659 | 15,688 ± 3,065 | 19.96 |
| AA1-s2 | A/A | TwinsUK | 74 | 3,040,994 | 17,011 ± 3,318 | 16.67 |
| AA2-t1 | A/A | TwinsUK | 24 | 5,936,129 | 16,830 ± 3,713 | 31.34 |
| AA2-t2 | A/A | TwinsUK | 36 | 5,353,906 | 14,053 ± 3,806 | 24.25 |
| AA3 | A/A | TwinsUK | 74 | 7,185,241 | 13,465 ± 2.589 | 31.19 |
| AA4 | A/A | TwinsUK | 47 | 6,065,965 | 11,014 ± 6,058 | 21.53 |
| AA5 | A/A | SL | 62 | 9,181,560 | 16,642 ± 4,891 | 53.64 |
| AA6 | A/A | SL | 31 | 8,634,738 | 17,226 ± 5,180 | 51.76 |
| AA7 | A/A | SL | 32 | 10,220,204 | 15,983 ± 4,614 | 57.04 |
| AA8 | A/A | SL | 31 | 8,876,456 | 16,081 ± 4,942 | 49.81 |
| AA9 | A/A | SL | 28 | 9,037,595 | 15,286 ± 4,748 | 48.79 |
| AN-s1 | A/new | TwinsUK | 27 | 6,113,622 | 15,827 ± 3,761 | 31.19 |
| AN-s2 | A/new | TwinsUK | 40 | 6,168,195 | 12,234 ± 4,412 | 24.32 |
| AB | A/B | SL | 33 | 6,982,155 | 16,533 ± 4,359 | 40.30 |
| AD | A/D | TwinsUK | 26 | 4,741,878 | 15,500 ± 3,477 | 23.69 |
| C1 | A/A | Blood | 0 | 12,659,998 | 16,632.1 ± 3,743 | 70.18 |
| C2 | A/A | Blood | 82 | 4,949,180 | 18,270.6 ± 1,753 | 30.12 |

**Table S1**: Sample pairs AA1-s1/AA1-s2 and AN-s1/AN-s2 are from the same donor at different ages. Samples AA2-t1 and AA2-t2 are monozygotic twins. C1 and C2 are negative control blood samples.

#### Supplementary Note on NCO tract length inference

We give a full likelihood model, where the observed data are the SNP locations and haplotype assignments for each read. We model the NCO tract length as a mixture of two geometric distributions. Our model assumes only simple CO and NCOs. Accordingly we only use reads with zero, one or two switches and discard "complex" reads (a switch being adjacent SNPs consistent with different haplotypes). The parameters we optimise are $q$, the probability of conversion of a recombination event to a CO; $m$, the probability of choosing the first out of the mixture of two geometric distributions of NCO tract lengths; and $l_1$ and $l_2$, the means of those two distributions. Let $\lambda_1 = 1/l_1$ and $\lambda_1 = 1/l_2$. Together we seek to find the MLE of

$$\theta = (q, m, l_1, l_2).$$

For a read with $n$ SNPs, denote by $p_1, \ldots, p_n$ the SNP positions along the read in bps; by $L$ the read length in bps; by $r_0$ the probability of a CO between the beginning of the read to the first SNP; by $r_1, \ldots, r_{n-1}$ the CO probability between each two adjacent SNPs, and by $r_n$ the probability of a CO between the last SNP to the end of the read. Additionally, we define $r_{-1}$ as the probability of a CO in a region (arbitrarily set to be $R = 5$kbp long) before the start of the read, which we will need later. Similarly, define $r_{n+1}$ as the probability of a CO in the region of length $R$ after read end. The $r_i$ quantities are obtained from a LD-based population-wide recombination map.

Our model also includes the assumption that a NCO tract is generated downstream, so we therefore have to use the two possible directions to take this assymetry into account. We do so by calculating the likelihood per read twice, once in the original direction and with the read positions and recombination probabilities in reverse order, and averaging the two. In the following we thus only describe the likelihood for the forward strand. We first describe each case separately.

**Two switches**: Two switches are only consistent with a NCO; we neglect the possibility of two COs within the same read. Suppose the first switch is between positions $i$ and $i+1$ and the second between $j$ and $j+1$. Then our likelihood is:

$$L_2 := \sum_{x=p_i}^{p_{i+1}-1} \sum_{y=p_j}^{p_{j+1}-1} \text{Pr(NCO starts at position x)} \cdot \text{Pr(NCO ends at position y|NCO starts at position x)}$$

The probability of a recombination event initiating and converting to a NCO between positions $i$ and $i+1$ is $r_i(1-q)/q$, so for any specific $x$ this is:

$$Pr(\text{NCO starts at position x}) = r_i(1-q)/((p_{i+1} - p_i)q)$$

The probability of finishing at $y$ given we have started at $x$ is the probability that a tract is of length $y-x$, which, according to the mixture of geometric distributions is:

$$\Pr(\text{NCO ends at position y}|\text{NCO starts at position x}) = m\cdot(1-\lambda_1)^{(y-x-1)}\lambda_1 + (1-m)\cdot(1-\lambda_2)^{(y-x-1)}\lambda_2$$

Together we get:

$$L_2 := \sum_{x=p_i}^{p_{i+1}-1}\sum_{y=p_j}^{p_{j+1}-1} r_i(1-q)/((p_{i+1}-p_i)q)\cdot\Big(m\cdot(1-\lambda_1)^{(y-x-1)}\cdot\lambda_1 + (1-m)\cdot(1-\lambda_2)^{(y-x-1)}\cdot\lambda_2\Big)$$

To simplify, we use the identity

$$\begin{aligned}
f_1(\lambda,A,B,C,D) :=& \sum_{x=A}^{B-1}\sum_{y=C}^{D-1}(1-\lambda)^{y-x-1}\cdot\lambda\\
=& \frac{\big((1-\lambda)^{-B}-(1-\lambda)^{-A}\big)\cdot\big((1-\lambda)^{C}-(1-\lambda)^{D}\big)}{\lambda}\\
=& \frac{1}{\lambda}\cdot(1-\lambda)^{-B}(1-(1-\lambda)^{B-A})\cdot(1-\lambda)^{C}(1-(1-\lambda)^{D-C})\\
=& \frac{1}{\lambda}\cdot(1-\lambda)^{C-B}\cdot(1-(1-\lambda)^{B-A})\cdot(1-(1-\lambda)^{D-C})
\end{aligned}$$

to get

$$L_2 = \frac{r_i(1-q)}{(p_{i+1}-p_i)q}\cdot\left(m\cdot f_1(\lambda_1,p_i,p_{i+1},p_j,p_{j+1}) + (1-m)\cdot f_1(\lambda_2,p_i,p_{i+1},p_j,p_{j+1})\right).$$

We further denote

$$f_2(A,B,C,D) := m\cdot f_1(\lambda_1,A,B,C,D) + (1-m)\cdot f_1(\lambda_2,A,B,C,D)$$

We can then write succinctly

$$L_2 = \frac{r_i(1-q)}{(p_{i+1}-p_i)q}\cdot f_2(p_i,p_{i+1},p_j,p_{j+1}).$$

**One switch.** One switch is consistent with either (i) a CO; (ii) a NCO where the first transition is unobserved and occured before the first SNP; or (iii) a NCO where the second transition is unobserved and occured after the last SNP. Suppose the switch occured between positions $i$ and $i+1$. For case (i), the probability of a CO between them is simply:

$$L_1^{CO} = r_i$$

For case (ii), the probability of a NCO starting at each position between the read start and $p_1$ is $r_0(1-q)/(p_1 q)$; and so as before, it is:

$$\frac{r_0(1-q)}{p_1 q} \cdot f_2(0, p_1, p_i, p_{i+1}).$$

Similarly, the probability of a NCO starting in the $R$ bp region before the read start and ending between positions $i$ and $i+1$ is:

$$\frac{r_{-1}(1-q)}{Rq} \cdot f_2(-R, 0, p_i, p_{i+1})$$

to get:

$$L_1^{NCO-left} = \frac{r_0(1-q)}{p_1 q} \cdot f_2(0, p_1, p_i, p_{i+1}) + \frac{r_{-1}(1-q)}{Rq} \cdot f_2(-R, 0, p_i, p_{i+1})$$

For case (iii), the probability that a geometric variable obtained a value above or equal to $t$ is $(1-\lambda)^{t-1}$. We will use the identity

$$\begin{aligned} f_3(\lambda, A, B, C) &:= \sum_{x=A}^{B-1} (1-\lambda)^{(C-x-1)} \\ &= \frac{(1-\lambda)^{C-B} - (1-\lambda)^{C-A}}{\lambda} \\ &= \frac{1}{\lambda} \cdot (1-\lambda)^{C-B}(1-(1-\lambda)^{B-A}). \end{aligned}$$

Similary define

$$f_4(A, B, C) := m \cdot f_3(\lambda_1, A, B, C) + (1-m) \cdot f_3(\lambda_2, A, B, C)$$

to get

$$L_1^{NCO-right} = \frac{r_i(1-q)}{(p_{i+1}-p_i)q} \cdot f_4(p_i, p_{i+1}, p_n).$$

Joining together we get:

$$L_1 := L_1^{CO} + L_1^{NCO-left} + L_1^{NCO-right}$$

**No switches.** A read with no switches is consistent with either: (i) no recominbation of any type; (ii) a CO before the first SNP or after the last SNP; (iii) a NCO tract contained between $R$ bp before read start and first SNP; (iv) a NCO tract contained between the last SNP and $R$ bp after read end; (v) a NCO tract contained between two adjacent SNPs; (vi) a NCO tract starting before the first SNP and ending after the last SNP. For case (i) the probability of no recombination is

$$L_0^{No} = 1 - \sum_{i=-1}^{n+1} r_i/q$$

where we neglect the possibility of two recombination events in the same read. For case (ii) the probability of a CO before the first SNP or after the last SNP is

$$L_0^{CO} = r_{-1} + r_0 + r_n + r_{n+1}$$

We next use the identity

$$f_5(\lambda, A, B) := \sum_{x=A}^{B-1} \sum_{y=x+1}^{B-1} (1-\lambda)^{y-x-1}\lambda = (B-A) - \frac{1-(1-\lambda)^{B-A}}{\lambda}$$

and define similarly

$$f_6(A, B) = m \cdot f_5(\lambda_1, A, B) + (1-m) \cdot f_5(\lambda_2, A, B)\,.$$

For case (iii), we have:

$$L_0^{NCO-left} = \frac{r_{-1}(1-q)}{R \cdot q} \cdot (f_6(-R, 0) + f_2(-R, 0, 0, p_1)) + \frac{r_0(1-q)}{p_1 \cdot q} \cdot f_6(0, p_1)\,.$$

For case (iv):

$$L_0^{NCO-right} = \frac{r_n(1-q)}{(L-p_n) \cdot q} \cdot (f_6(p_n, L) + f_4(p_n, L, L)) + \frac{r_{n+1}(1-q)}{R \cdot q} \cdot (f_6(L, L+R) + f_4(L, L+R, L+R))\,.$$

For case (v), we define the probability of an NCO between each pair of SNPs as:

$$L_0^{NCO-between} = \frac{r_1(1-q)}{(p_2-p_1)q} \cdot f_6(p_1, p_2) + \ldots + \frac{r_{n-1}(1-q)}{(p_n-p_{n-1})q} \cdot f_6(p_{n-1}, p_n)$$

For case (vi):

$$L_0^{NCO-all} = \frac{r_{-1}(1-q)}{R \cdot q} \cdot f_4(-R, 0, p_n) + \frac{r_0(1-q)}{p_1 \cdot q} \cdot f_4(0, p_1, p_n)$$

Finally we define

$$L_0 := L_0^{No} + L_0^{CO} + L_0^{NCO-left} + L_0^{NCO-right} + L_0^{NCO-between} + L_0^{NCO-all}$$

**Total likelihood.** The total likelihood for the forward direction is

$$L^{forward} := L_0 + L_1 + L_2$$

and the final total likelihood is

$$L := (L^{forward} + L^{backward})/2$$

with $L^{backward}$ calculated the same $L^{forward}$ but with the read and CO probabilities reversed. The likelihood of a set of reads is the product of the likelihoods per read. We used the Nelder-Mead algorithm to optimize.
